# Homeostatic control of stem cell activity during intestinal regeneration

**DOI:** 10.64898/2026.07.15.738595

**Authors:** Shichao Yang, Changsheng Luo, Guofan Peng, Kewei Zheng, Kang Han, Jun Zhou

## Abstract

Stem cells proliferate rapidly to maintain fast tissue turnover during regeneration. However, the feedback mechanisms in stem cells that prevent hyperproliferation remain unclear, and their dysregulation can lead to organ failure and cancer. Here, we identified nuclear factor-Y (NF-Y) as the transcriptional repressors to maintain the stem cell quiescence during intestinal homeostasis. We found that NF-Y negatively regulates intestinal stem cell (ISC) proliferation through preferentially occupying the promoters of EGFR signaling pathway components Egfr/Mkp3/Raf/Ras/pointed, via the action of histone acetyltransferase Nejire (Nej)/p300 dependent transcription regulation. While the loss of NF-Y enhances ISC proliferation, cell death and sensitivity to stress and tumor induced mortality. Moreover, NF-Y acts together with Nej to restrict Egfr expression and suppress ISC hyperproliferation. Together, these results demonstrate NF-Y acts with Nej serve as a key negative feedback module to orchestrate transcription initiation and termination of growth signaling in the control of stem cell activity in homeostatic and disease conditions.

## Introduction

The ability of adult tissues to maintain homeostasis and mount regenerative responses following injury is fundamentally dependent on the activity of tissue-resident stem cells (1). Among the various model systems available for studying stem cell biology, the Drosophila melanogaster midgut has emerged as a particularly powerful and tractable system. The fly midgut has remarkable structural, functional, and genetic conservation with the mammalian intestine, combined with sophisticated genetic tools, allows for detailed mechanistic dissection of stem cell regulation at single-cell resolution within a living organism (2, 3). The adult Drosophila midgut is a highly dynamic epithelial tissue that undergoes continuous renewal throughout the organism’s lifespan. This renewal is driven by a population of multipotent intestinal stem cells (ISCs) that are interspersed along the basal side of the epithelial monolayer. ISC is capable of producing its differentiated progeny, creating a stereotyped cellular architecture that facilitates lineage tracing and functional analyses. Enteroblasts subsequently differentiate into either absorptive enterocytes, which constitute the majority of the epithelial lining and are responsible for nutrient absorption, or secretory enteroendocrine cells, which regulate gut physiology to maintain tissue homeostasis (4–7). This carefully balanced lineage output ensures that the cellular composition of the intestinal epithelium remains stable despite constant turnover.

Precise control of ISC proliferation, differentiation, and quiescence is essential for maintaining tissue homeostasis. Disruption of this balance can have severe consequences. For instance, excessive proliferation leads to hyperplasia and can predispose to tumor formation, while insufficient proliferation compromises epithelial integrity and barrier function, leading to increased susceptibility to infection and premature aging (8–12). Under normal conditions, ISCs exhibit relatively low proliferative activity, entering the cell cycle only when needed to replace aged or damaged cells. This quiescent state is actively maintained by multiple intrinsic and extrinsic regulatory mechanisms that prevent inappropriate stem cell activation. The regenerative capacity of the intestinal epithelium becomes particularly evident upon tissue damage. The Drosophila midgut is continuously exposed to environmental insults, including ingested pathogens, oxidative stress, and mechanical injury. In response to such challenges, ISCs rapidly exit quiescence and enter a state of heightened proliferative activity to replenish lost or damaged cells (13–15). Upon tissue damage, damaged enterocytes release mitogenic signals, including ligands of the Wnt/Wingless, InR, JAK-STAT and Toll pathways, which activate ISCs in a paracrine manner (8, 15–20). Epidermal growth factor receptor (EGFR) signaling is a principal driver of intestinal stem cell (ISC) proliferation and regeneration in the Drosophila midgut (14, 18, 21). Following tissue damage, damaged enterocytes secrete EGFR ligands, which activate the Egfr receptor and its downstream signaling cascade on ISCs to promote their rapid division and replenishment of lost cells(8, 14). Conversely, excessive EGFR activation leads to ISC hyperproliferation and tumor-like overgrowth, highlighting the necessity of tight negative regulation to maintain homeostasis (14, 22).

Following the resolution of damage, the proliferative response must be terminated to prevent excessive growth and tissue overgrowth. Failure to re-establish stem cell quiescence leads to persistent hyperplasia, which can progress to dysplastic lesions and tumor-like structures (11, 12, 22–24). This transition from active proliferation back to quiescence requires robust negative feedback mechanisms that sense when tissue integrity has been restored and subsequently suppress mitogenic signaling. While numerous studies have elucidated the pro-proliferative signals that activate ISCs during regeneration, the molecular mechanisms that enforce the cessation of proliferation and prevent hyperproliferation remain comparatively less understood. Identifying these negative regulators is critical, as their dysfunction is increasingly recognized as a contributing factor to inflammatory bowel diseases, age-related decline in regenerative capacity, and intestinal cancers (16, 17, 25).

Transcriptional regulation plays a central role in controlling stem cell behavior, governing the expression of genes that drive proliferation, differentiation, and quiescence. Nuclear Factor Y (NF-Y) is a highly conserved heterotrimeric transcription factor complex present in all eukaryotes(26, 27). It consists of three subunits: NF-YA, NF-YB, and NF-YC. NF-YB and NF-YC each contain a histone-fold domain (HFD), which enables them to form a stable heterodimer in the cytoplasm before translocating to the nucleus, where they recruit NF-YA to assemble the functional heterotrimer(28). The NF-YA subunit confers sequence-specific DNA binding to CCAAT-box motifs in gene promoters, while the NF-YB/NF-YC dimer structurally resembles the histone H2A/H2B dimer and facilitates chromatin interaction(27, 28). Beyond acting as a transcriptional activator for many cell cycle and housekeeping genes, NF-Y can also function as a transcriptional repressor in specific contexts by recruiting co-repressors or modulating chromatin accessibility(27, 29). In *Drosophila*, NF-Y plays important roles in development, stress response, and tissue homeostasis(30, 31), although its function in adult stem cell regulation remains largely unexplored. In this study, we investigate the role of NF-Y as a transcriptional repressor that maintains ISC quiescence during intestinal homeostasis and functions as a key negative feedback module during tissue damage and repair. Our findings reveal that NF-Y, acting together with the histone acetyltransferase Nejire/p300, coordinates the transcriptional repression of key growth signaling components, including Egfr, Mkp3, Raf, Ras, and Pnt, thereby ensuring the timely termination of ISC proliferation following regenerative episodes. Elucidating this mechanism not only advances our understanding of intestinal stem cell biology but also are likely applicable to other tissue stem cells in regenerative conditions and diseases.

## Results

### The NF-Y complex is a stem cell regulator in intestinal regeneration

To identify transcriptional regulators of intestinal stem cell (ISC) activity during gut regeneration, we established a Drosophila model of intestinal injury and recovery using dextran sodium sulfate (DSS)(32, 33). Feeding adult flies 3% DSS induced robust epithelial damage, as evidenced by a marked increase in mitotic activity quantified by phospho-histone H3 (PH3) staining. Notably, PH3 levels exhibited two distinct peaks at day 2 (D2) and day 6 (D6) during the injury and recovery time course, reflecting successive waves of regenerative proliferation corresponding to the acute damage response and subsequent tissue repair (Fig. 1 *A*). This biphasic mitotic pattern provided a defined temporal framework for downstream analyses of molecular signatures. We therefore performed bulk RNA-seq on dissected midguts collected at four time points: D1 and D4 during DSS treatment, and recovery day 1 (corresponding to D5) and recovery day 4 (corresponding to D8) after transfer to a normal diet (Fig. 1 *B*). Principal component analysis (PCA) revealed that control, DSS treated, and recovery samples occupied distinct transcriptional states, with PC1 (31.62%) and PC2 (21.33%) together capturing the major axes of variation (Fig. 1 *C*). These data indicate that intestinal injury and repair are accompanied by large-scale, temporally ordered transcriptional reprogramming.

**Figure 1.**
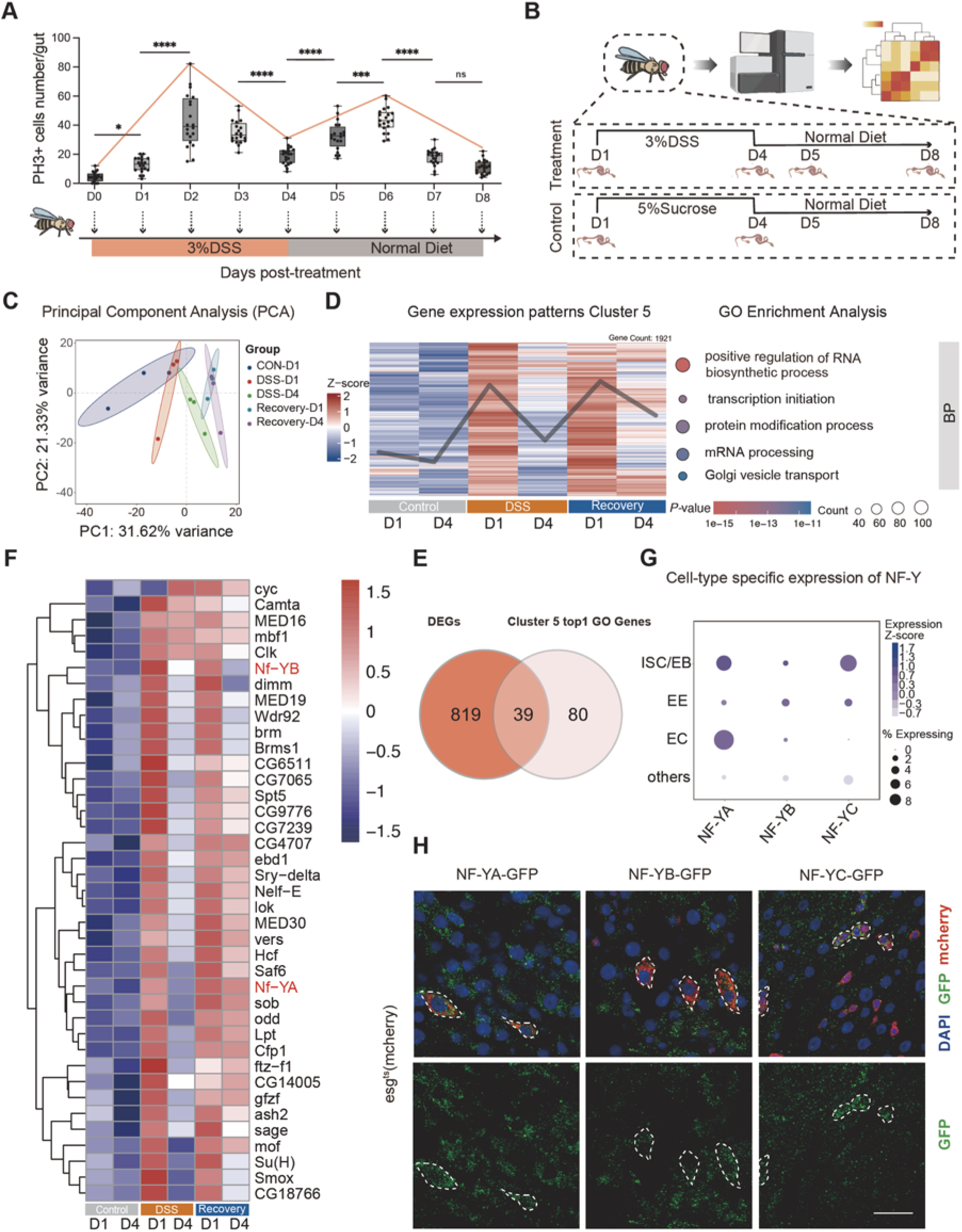
Integrative transcriptomic analysis identifies the NF-Y complex as a regulator of intestinal regeneration. (A) Quantification of PH3+ mitotic cells per gut at indicated time points during 3% DSS induced injury (pink bar) and subsequent recovery on normal diet (gray bar). Data are presented as box- and whisker plots, where boxes extend from the 25th to 75th percentiles, the horizontal line indicates the median, and whiskers represent the minimum and maximum values. Each dot represents an individual gut (n ≥ 20 guts per time point). ****p < 0.0001, ***p < 0.001; analyzed by one-way ANOVA with Sidak’s multiple comparisons test. (B) Schematic view of the bulk RNA-seq experimental design. (C) PCA of midgut transcriptomes across control, DSS treated, and recovery conditions. (D) Temporal gene expression clustering during intestinal regeneration. (Left) Heatmap of Cluster 5 genes (n = 1921) showing progressive upregulation during injury and sustained elevation through recovery. (E) Venn diagram illustrating the overlap between all DEGs identified and Cluster 5 “positive regulation of RNA biosynthetic process” term genes. (F) Heatmap of 39 DEGs across control (D1, D4), DSS (D1, D4), and recovery (D1, D4) stages. NF-YA and NF-YB exhibit coordinated dynamic upregulation aligned with the regenerative response. (G) Cell type specific expression of NF-Y subunits in the adult midgut. Dot plot derived from published single-cell RNA-seq data showing expression levels. (H) NF-Y subunits are expressed in intestinal stem/progenitor cells. Representative images of adult midguts from flies expressing esg-Gal4 > UAS-mCherry (red) carrying endogenously tagged NF-YA-GFP, NF-YB-GFP, or NF-YC-GFP (green). Nuclei are stained with DAPI (blue). Dashed lines demarcate esg+ cell clusters. Scale bar, 50 μm.

To identify genes dynamically regulated during regeneration, we performed weighted gene co expression clustering across all sampled time points and defined eight distinct temporal clusters C1-C8 (Fig. S1 *A*). Among these, Cluster 5 (n = 1,921 genes) exhibited a similar biphasic expression pattern (Fig. 1 *D*, left). Gene Ontology (GO) analysis of Cluster 5 revealed strong enrichment for processes related to RNA biosynthesis and transcriptional initiation (Fig. 1 *D*, right), suggesting that this cluster captures transcriptional programs engaged at multiple stages of the regenerative response. In parallel, Cluster 3 (n = 1,624 genes) showed a temporal pattern opposite to that of Cluster 5, with a sharp and transient peak at late injury (DSS D4) and rapid decline during recovery (Fig. S1 *B*). This cluster was enriched for processes including protein refolding, chromosome organization, and cell projection assembly, indicating distinct functional waves of gene expression during regeneration.

We next prioritized high confidence regulatory candidates by intersecting significantly upregulated genes at DSS D1 with genes associated with the top ranked GO term, “positive regulation of RNA biosynthetic process,” enriched in Cluster 5. This analysis identified 39 overlapping genes whose expression was tightly coupled to the regenerative response (Fig. 1 *E*). Among these candidates, a heatmap of transcription-related DEGs revealed that two subunits of the Nuclear Factor Y (NF-Y) complex, NF-YA and NF-YB, exhibited strong, coordinated dynamic upregulation across the injury and recovery stages (Fig. 1 *F* and Fig. S1 *C*). In contrast, the expression of NF-YC, remained relatively stable throughout the regenerative process. However, basal transcriptomic quantification demonstrated that NF-YC is constitutively expressed at substantially higher levels than NF-YA and NF-YB in the homeostatic midgut (Fig. S1 *D*). Given this baseline abundance, the dynamic stress-induced upregulation of the rate-limiting NF-YA and NF-YB subunits is sufficient to drive the functional assembly and activation of the entire NF-Y complex. Together, these observations nominate the NF-Y complex as a critical transcriptional regulator of intestinal regeneration.

We then examined the cellular distribution of NF-Y subunits in the adult midgut. Analyzing published single cell RNA-seq data (34), we found that NF-YA, NF-YB, and NF-YC were most highly expressed in ISC/enteroblast (EB) populations relative to enteroendocrine cells (EEs) and enterocytes (ECs) (Fig. 1 *G*). Using endogenously GFP tagged NF-YA, NF-YB, and NF-YC transgenes, we confirmed that all three subunits were presented in esg+ stem/progenitor cells under homeostatic conditions (Fig. 1 *H*).

To determine whether NF-Y induction reflects a broader stress response, we examined NF-Y expression in additional intestinal injury contexts. NF-YA, NF-YB, and NF-YC were all induced following oral infection with the enteric pathogen *Pseudomonas entomophila* (35), with upregulation evident at both 4 h and 16 h after infection (Fig. S1 *E*). NF-Y expression was also markedly elevated in midguts with Notch depletion, a genetic model of ISC hyperproliferation and tumor like overgrowth (36) (Fig. S1 *F*). These findings indicate that induction of the NF-Y complex is a common transcriptional response to different intestinal stress.

Finally, we examined the relevance of NF-Y to human disease. Analysis of GEPIA2 expression data, based on TCGA and GTEx datasets, revealed elevated expression of *NFYA* in both colon adenocarcinoma (COAD) and rectal adenocarcinoma (READ) relative to normal tissues (Fig. S1 *G*)(37). Moreover, survival analysis using Kaplan–Meier Plotter showed that high *NFYA* expression was associated with significantly better overall survival in colorectal cancer patients (HR = 0.64, 95% CI: 0.51-0.82; log-rank p = 0.00029) (Fig. S1 *H*)(38). Together, these multi layered analyses identify the NF-Y complex as a dynamically regulated transcriptional regulator associated with intestinal regeneration in Drosophila and colorectal cancer in humans.

### The NF-Y complex inhibits intestinal stem cell expansion and maintains homeostasis

We next examined the protein expression dynamics of NF-Y complex and tested its function in vivo. Endogenously tagged NF-YA-GFP was monitored in esg+ progenitor cells across the DSS injury and recovery time course. Consistent with the transcriptomic data, NF-YA protein levels increased during DSS treatment, peaked at late injury, and declined toward baseline during recovery. Notably, this dynamic expression pattern was observed not only at the whole cell level but also specifically within the nuclei of progenitor cells (Fig. 2 *A-B* and Fig. S2 *B*). This dynamic regulation in the progenitor compartment suggested that NF-Y may function directly during the regenerative response.

**Figure 2.**
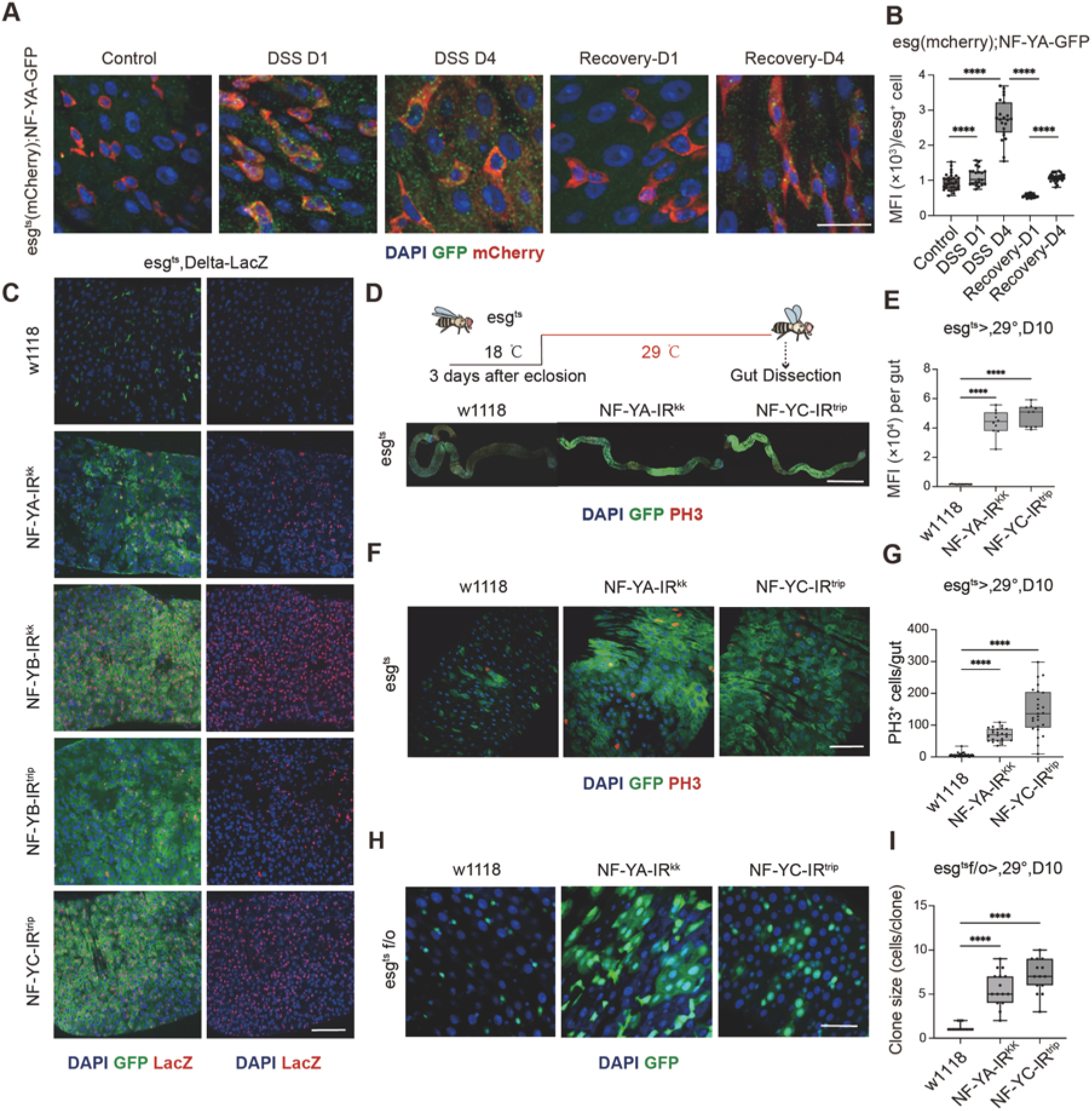
The NF-Y complex is required to maintain intestinal homeostasis (A and B) Dynamic expression of NF-YA in intestinal progenitor cells during regeneration. (A) Representative confocal images of adult midguts expressing esg-Gal4 > UAS-mCherry (red) and endogenously tagged NF-YA-GFP (green) under homeostatic, DSS injured and recovery conditions. Nuclei are stained with DAPI (blue). Scale bar, 20 μm. (B) Quantification of NF-YA-GFP mean fluorescence intensity (MFI) within esg+ progenitor cells at indicated time points. Data are presented as box-and-whisker plots. *p < 0.05, ****p < 0.0001; analyzed by one-way ANOVA with Sidak’s multiple comparisons test (n ≥ 20 guts per time point). (C) Loss of NF-Y subunits induce expansion of the ISC pool. Representative images of midguts from control (w1118) and NF-Y knockdown (NF-YA-IR, NF-YB-IR, NF-YC-IR) flies. Delta-LacZ (red) marks ISCs and esg^ts^ > GFP (green) marks ISC/EB populations. Scale bar, 50 μm. (D and E) Global expansion of progenitor cells upon NF-Y depletion. (D) Schematic view of the experiment plan (top) and representative images of whole midguts (bottom) showing the distribution of esg+ cells (green) and PH3+ mitotic cells (red). Scale bar, 1 mm. (E) Quantification of total GFP MFI per gut, reflecting overall progenitor cell expansion. n ≥10 guts per group. (F and G) NF-Y depletion promotes ISC hyperproliferation during normal homeostasis. (F) representative image of midguts expressing *esg^ts^ > GFP* (green) stained for the mitotic marker PH3 (red). (G) Quantification of PH3+ cells per gut. n ≥24 guts per group. (H and I) NF-Y restricts ISC lineage expansion. (H) Representative confocal images of lineage tracing using the esg^ts^ flip/out system (esg^ts^ F/O) in control and NF-Y knockdown midguts at 10 days post induction. GFP (green) marks progeny derived from esg+ progenitor cells. Scale bar, 50 μm. (I) Quantification of clone size (cells per clone), showing accelerated lineage expansion upon NF-Y depletion. n ≥15 guts per group. (Statistical note for E, G, I: Data are presented as box and whisker plots indicating the median, 25th – 75th percentiles, and minimum/maximum values. analyzed by one-way ANO VA with Dunnett’s multiple comparisons test; *P < 0.05, **P < 0.01, ***P < 0.001, ****P < 0.0001).

To validate the efficiency of the NF-Y knockdown lines, we first confirmed that expression of NF-YA, NF-YB, and NF-YC was significantly reduced in the respective RNAi lines (Fig. S2 *A*).We then knocked down NF-YA, NF-YB, or NF-YC in the midgut using esg^ts^ driver combined with stem cell reporter Delta-lacZ, and noted a marked expansion of the progenitor compartment, characterized by increased numbers and density of both esg>GFP+ cells and Delta-LacZ+ ISC enriched cells (Fig. 2 *C*)(39, 40). This phenotype was evident across broad regions of the midgut and resulted in a significant increase in total progenitor cell burden (Fig. 2 *D* and *E*). These data indicate that the NF-Y complex is required to constrain the expansion of the intestinal progenitor compartment during homeostasis.

As the expansion of the esg+ compartment suggested elevated proliferative activity, we next assessed mitosis by PH3 staining and found a strong increase in dividing cells in midguts depleted of NF-YA or NF-YC relative to controls (Fig. 2 *F* and *G*). To evaluate this hyperproliferative state as stem cell lineage expansion, we performed lineage tracing analysis using the esg^ts^ flip/out system (41). After 7 days of induction, clones generated in NF-YA or NF-YC depleted intestines were significantly larger than those in control guts, indicating increased stem cell proliferation and differentiation (Fig. 2 *H* and *I*). Together, these results show that NF-Y restrains ISC proliferation and lineage expansion to maintain intestinal homeostasis.

To determine whether the NF-Y depletion induced hyperproliferation is restricted in ISC, we next depleted NF-YA and NF-YC in differentiated enterocytes using Myo1Ats(42, 43). In contrast to progenitor specific knockdown, EC specific depletion had no detectable effect on ISC proliferation (Fig. S2 *C* and *D*), indicating that NF-Y mediated stem cell function is intrinsic to the ISC/EB compartment. We further validated this using a temporal CRISPR/Cas9 strategy in esg+ progenitor cells. The NF-Y knockout mutants showed sustained ISC hyperproliferation, phenocopying the RNAi based results (Fig. S2 *E-G*). These findings indicate that NF-Y is required within the progenitor compartment to maintain long term proliferative homeostasis. Collectively, these data identify the NF-Y complex as a key regulator to control ISC quiescence in the adult midgut.

### NF-Y complex constricts intestinal regeneration in both stress and tumor conditions

We next asked whether loss of NF-Y would further compromise the intestinal response to epithelial injury. To address this, flies expressing NF-YA or NF-YC RNAi in esg+ intestinal progenitor cells were first maintained on a normal diet at 29°C for 7 days to ensure efficient NF-Y depletion, and were then challenged with 3% DSS for 3 additional days before analysis (Fig. 3 *A*)(33, 44). Under these injury conditions, NF-Y depleted midguts displayed a marked increase in PH3 positive mitotic cells compared with controls (Fig. 3 *B* and *C*), indicating that NF-Y is required to limit ISC proliferation during tissue damage. Loss of NF-Y also increased the incidence and severity of the Smurf phenotype following DSS exposure (Fig. 3 *D*). a comparable short-gut phenotype was also observed when NF-Y-depleted intestines were subjected to DSS induced injury (Fig. S3 *A* and *B*). In addition, NF-YA and NF-YC depleted flies exhibited significantly reduced survival during DSS challenge (Fig. 3 *E*). These findings indicate that NF-Y complex acts as a brake to restrict ISC hyperproliferation, which is essential for maintaining barrier function and homeostasis during damage response (Fig. S3 *C*).

**Figure 3.**
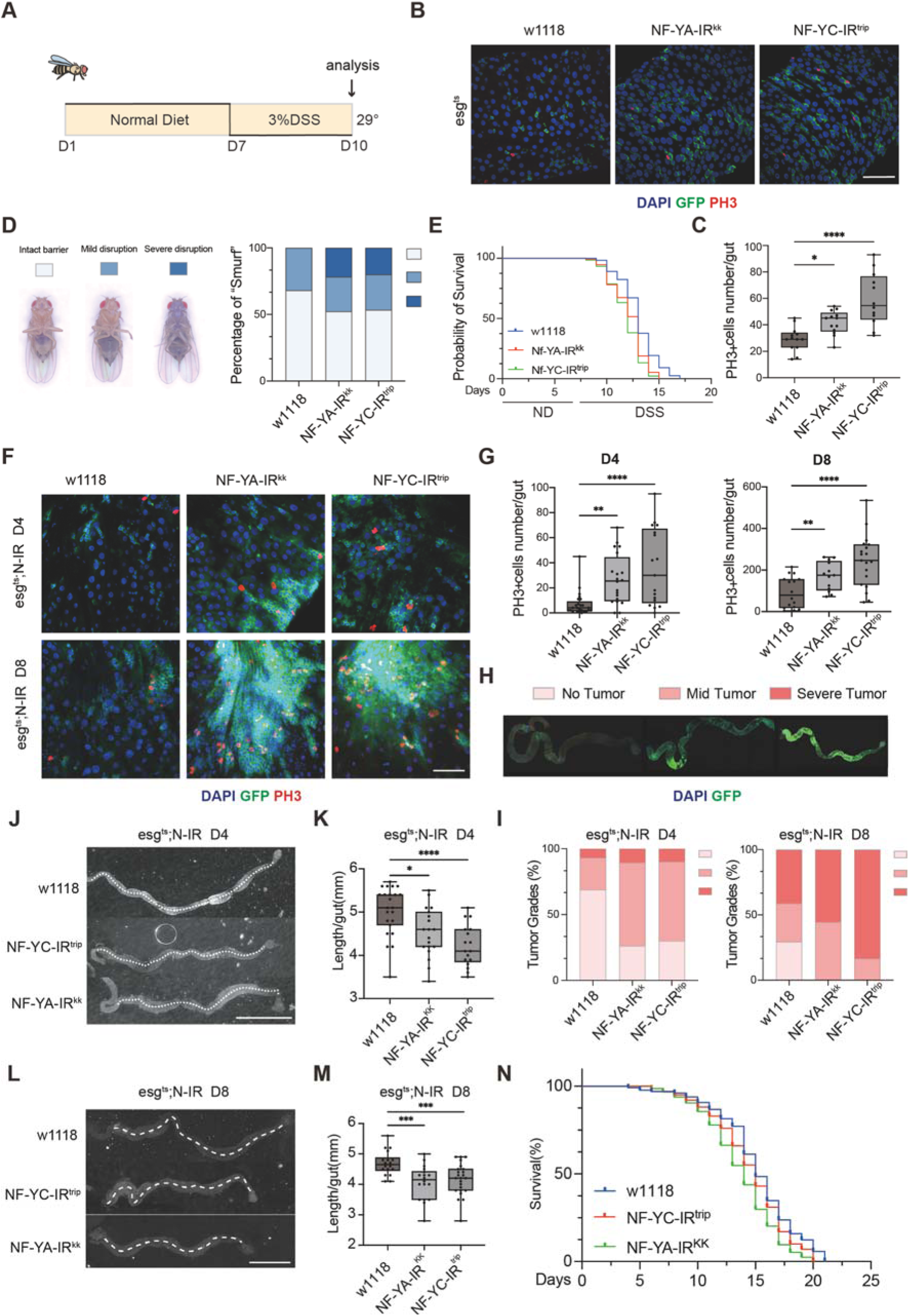
Loss of NF-Y exacerbates DSS induced injury and induce tumor like overgrowth. (A) Schematic view of the experiment plan. (B and C) Representative images (B) and quantification (C) of PH3⁺ mitotic cells in midguts from control (w1118) and NF-YA or NF-YC RNAi flies. NF-Y depletion increased mitotic activity under injury conditions. n ≥ 12 midguts per group; Blue, DAPI; green, GFP; red, PH3; scale bar, 50 µm. (D) Representative images of the Smurf phenotype and quantification of barrier disruption categories (intact, mild, severe) for each genotype after DSS treatment. NF-Y depleted intestines showed severe barrier defect. n ≥ 30 midguts per group. (E) Kaplan–Meier survival analysis for control and NF-Y depleted flies after DSS challenge. NF-Y knockdown significantly reduced survival during injury. n ≥ 100 flies per genotype; log rank (Mantel–Cox) test; ****P < 0.0001. (F and G) NF-Y depletion exacerbates hyperproliferation in the Notch knockdown (N-IR) tumor model. Representative images (F) and quantification (G) of PH3⁺ cells per midgut at D4 and D8 with NF-Y RNAi. Mitotic activity increased in NF-Y depleted intestines. n ≥ 17 midguts per group; Blue, DAPI; green, GFP; red, PH3; scale bar, 50 µm. (H and I) Representative images of whole gut (H) and quantification of tumor grades (I) at D4 and D8. Midguts were classified as no tumor, mild tumor or severe tumor. NF-Y knockdown shifted the distribution toward more severe tumor like phenotypes. n ≥ 30 midguts per group; Blue, DAPI; green, GFP; red, PH3. (J–M) Representative images of whole gut at D4 (J) and D8 (L), together with quantification of gut length at D4 (K) and D8 (M) in the N-IR background. White dashed lines indicate the measured longitudinal axis. NF-Y depletion resulted in progressive gut shortening and altered intestinal morphology. n ≥ 17 midguts per group; Blue, DAPI; green, GFP; red, PH3; Scale bar, 1 mm. (N) Kaplan–Meier survival curves for flies with NF-Y knockdown, showing shortened lifespan relative to controls. n ≥ 100 flies per genotype; log rank test; ****P < 0.0001. (Statistical note for C, G, K, M: Data are presented as box and whisker plots indicating the median, 25th–75th percentiles, and minimum/maximum values. analyzed by one-way ANOVA with Dunnett’s multiple comparisons test; *P < 0.05,**P < 0.01, ***P < 0.001, ****P < 0.0001).

We next sought to determine whether NF-Y mediated inhibition of ISC proliferation is essential under disease conditions, such as in tumors. We found that knockdown of NF-YA or NF-YC in notch depleted midgut display a significant increase in mitotic activity in both D4 and D8 (Fig. 3 *F* and *G*). Whole gut imaging further revealed an increased severity of dysplastic overgrowth in NF-Y deficient tumor bearing guts. Because the expansion of esg>GFP fluorescence directly reflects the intestinal stem cell burden,thereby delineating tumor size and location.we observed a clear shift from mild to severe lesion states relative to controls (Fig. 3 *H* and *I*). We further noted that midguts lacking NF-YA or NF-YC became visibly shortened and distorted, and quantitative analysis confirmed a significant reduction in gut length at both D4 and D8 relative to controls (Fig. 3 *J-M*). Loss of NF-Y additionally shortened overall lifespan (Fig. 3 *N*), Similar phenotype was observed in a Ras^V12^/Apc-IR tumor model (45, 46), NF-Y depletion leads to enhanced mitotic activity, advanced tumor grades, exacerbated gut shortening, and accelerated mortality (Fig. S3 *D-I*). These findings indicate that NF-Y complex is capaple of inhibiting intestinal tumor progression. Hence, these results identify NF-Y as a critical safeguard of intestinal barrier function to restrict ISC hyperproliferation in both stress and disease conditions.

### NF-Y complex influences transcriptional program to regulate ISC function

We next sought to investigate the molecular mechanism on how NF-Y controls ISC quiescence in the midgut. To this end, we performed RNA-seq on midguts from control and flies expressing NF-YA RNAi in esg+ progenitor cells after 10 days of induction at 29°C (Fig. 4 *A*). PCA analysis revealed clear separation between control and NF-YA depleted samples, indicating that loss of NF-YA induces substantial transcriptomic changes in the intestine (Fig. 4 *B*). Differential expression analysis identified widespread gene expression changes, including deregulation of multiple genes linked to proliferative control and signaling (Fig. 4 *C*). KEGG pathway analysis of differentially expressed genes highlighted pathways associated with tissue growth and cell fate regulation, including MAPK, Notch, and Hippo signaling (Fig. 4 *D*). Gene set enrichment analysis further confirmed significant enrichment of the MAPK signaling pathway upon NF-YA depletion (Fig. 4 *E*). GO analysis of the same RNA-seq comparison revealed enrichment for biological processes related to mitotic progression and cell division, while additional GSEA identified enrichment of the Toll and Imd signaling pathway (Fig. S4 *A* and *B*). Together, these data indicate that NF-Y depletion triggers broad transcriptional reprogramming linked to proliferative, stress responsive, and developmental signaling programs.

**Figure 4.**
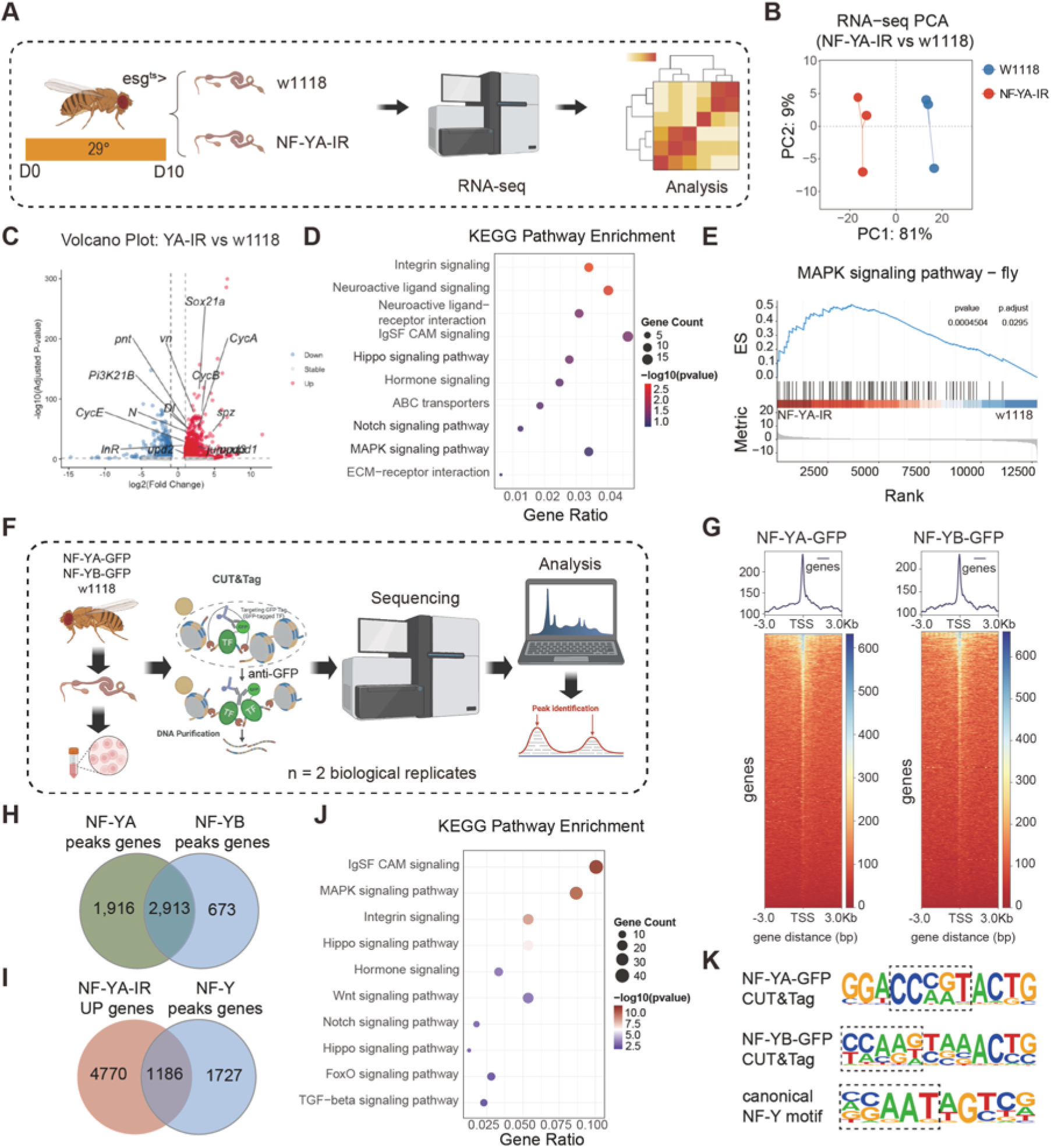
Integrated RNA-seq and CUT&Tag analyses reveal NF-Y influences transcriptional programs. (A) Schematic view of the RNA-seq experimental design. (B) Principal component analysis (PCA) of RNA-seq samples from control and NF-YA depleted midguts. (C) Volcano plot showing differentially expressed genes (DEGs) in NF-YA depleted versus control (w1118) midguts. Selected deregulated genes are highlighted. (D) KEGG pathway enrichment analysis of DEGs identified between NF-YA depleted and control (w1118) midguts. (E) GSEA showing significant enrichment of the MAPK signaling pathway in NF-YA depleted midguts. NES and adjusted p value are indicated in the plot. (F) Schematic view of the CUT&Tag workflow to profile target of NF-YA-GFP and NF-YB-GFP in the adult midgut. (G) Metagene plots and heatmaps showing NF-YA and NF-YB CUT&Tag signal centered on transcription start sites (TSSs), indicating promoter-proximal enrichment of NF-Y binding. (H) Venn diagram showing the overlap between NF-YA and NF-YB peak-associated genes. (I) Venn diagram showing the overlap between NF-YA depleted upregulated genes and NF-Y complex peak associated genes, identifying 1,186 NF-Y target genes. (J) KEGG pathway enrichment analysis of the NF-Y target genes shown in (I). These NF-Y targets are enriched in signaling pathways including MAPK, Wnt, Notch, Hippo, and TGF-β. (K) De novo motif analysis of NF-YA-GFP and NF-YB-GFP CUT&Tag peaks. The identified motifs closely match the canonical NF-Y-binding motif (CCAAT box).

As NF-Y is a conserved transcription factor complex, we next profiled NF-Y chromatin occupancy in the adult midgut using CUT&Tag analysis (Fig. 4 *F*)(47). CUT&Tag analysis revealed strong enrichment of both NF-YA and NF-YB signals around transcription start sites (TSSs), indicating that NF-Y binding is predominantly promoter proximal (Fig. 4 *G*). Motif enrichment analysis of NF-YA and NF-YB CUT&Tag peaks identified strong enrichment of the canonical NF-Y binding motif (CCAAT box) (Fig. 4 *K*). Genes associated with NF-YA and NF-YB peaks showed extensive overlap, defining a large shared set of candidate NF-Y bound target genes (Fig. 4 *H*).

We then integrated the RNA-seq and CUT&Tag datasets to identify candidate direct NF-Y target genes that are normally restrained by the NF-Y complex. Intersecting genes upregulated upon NF-YA depletion with NF-Y complex peak associated genes identified 1,186 overlapping genes (Fig. 4 *I*), representing a high confidence set of candidate direct NF-Y target genes. KEGG analysis of this overlapping gene set revealed significant enrichment for pathways involved in growth control and developmental regulation, including MAPK, Wnt, Notch, Hippo, FoxO, and TGF-β signaling (Fig. 4 *J*). GO analysis of the same 1186 gene set showed enrichment for morphogenetic and developmental processes, including tissue morphogenesis, epithelial cell differentiation, cell projection, and growth (Fig. S4 *E*). Overlap analysis further showed that genes associated with the EGFR pathways were substantially represented among NF-YA upregulated genes (Fig. S4 *C*), and heatmap analysis confirmed broad transcriptional upregulation of MAPK related genes in NF-YA depleted midguts (Fig. S4 *D*). Genome browser inspection further revealed prominent NF-YA and NF-YB occupancy at the Egfr and Mkp3 loci, together with promoter-associated H3K4me3 signal (48), consistent with direct NF-Y binding at MAPK associated target loci (Fig. S4 *F* and *G*). Furthermore, motif analysis specifically performed on the promoter regions of core EGFR signaling pathway genes also showed significant enrichment of the NF-Y binding motif (Fig. S4 *H*). Together, these integrated transcriptomic and chromatin profiling analyses identify the NF-Y complex as a direct repressor of growth and regeneration associated signaling programs.

### NF-Y complex acts with Nej to control stem cell activity

To explore how NF-Y regulates numerous major growth signaling pathways, we search for the Protein association network analysis and identified Nej, the *Drosophila* homolog of the histone acetyltransferase coactivator CBP/p300, as a candidate NF-Y associated regulatory factor. Building upon this computational prediction, we experimentally substantiated the physical interaction between NF-Y and Nej using co-immunoprecipitation (Co-IP) assays (Fig. 5 *A*). Previous studies in mammalian systems showing that NF-YA physically interact with CBP/p300 in sympathetic neurons to regulate NF-Y target expression (49–51). We next tested whether Nej contributes to the chromatin state changes upon NF-Y depletion. We therefore performed ATAC-seq on midguts from control, NF-YA RNAi, and flies co-expressing NF-YA RNAi and nej RNAi (Fig. 5 *B*). Metaplot and heatmap analyses revealed that NF-YA depletion caused a pronounced increase in chromatin accessibility around promoter-proximal regions, whereas co-depletion of nej largely attenuated this genomic accessibility (Fig. 5 *B*). Genome-wide visualization of open chromatin peaks and genomic feature annotation further supported the overall quality of the ATAC-seq datasets and their promoter-enriched distribution (Fig. S5 *A* and *B*). These findings indicate that the increase chromatin accessibility upon NF-Y depletion depends, at least in part, on Nej activity.

**Figure 5.**
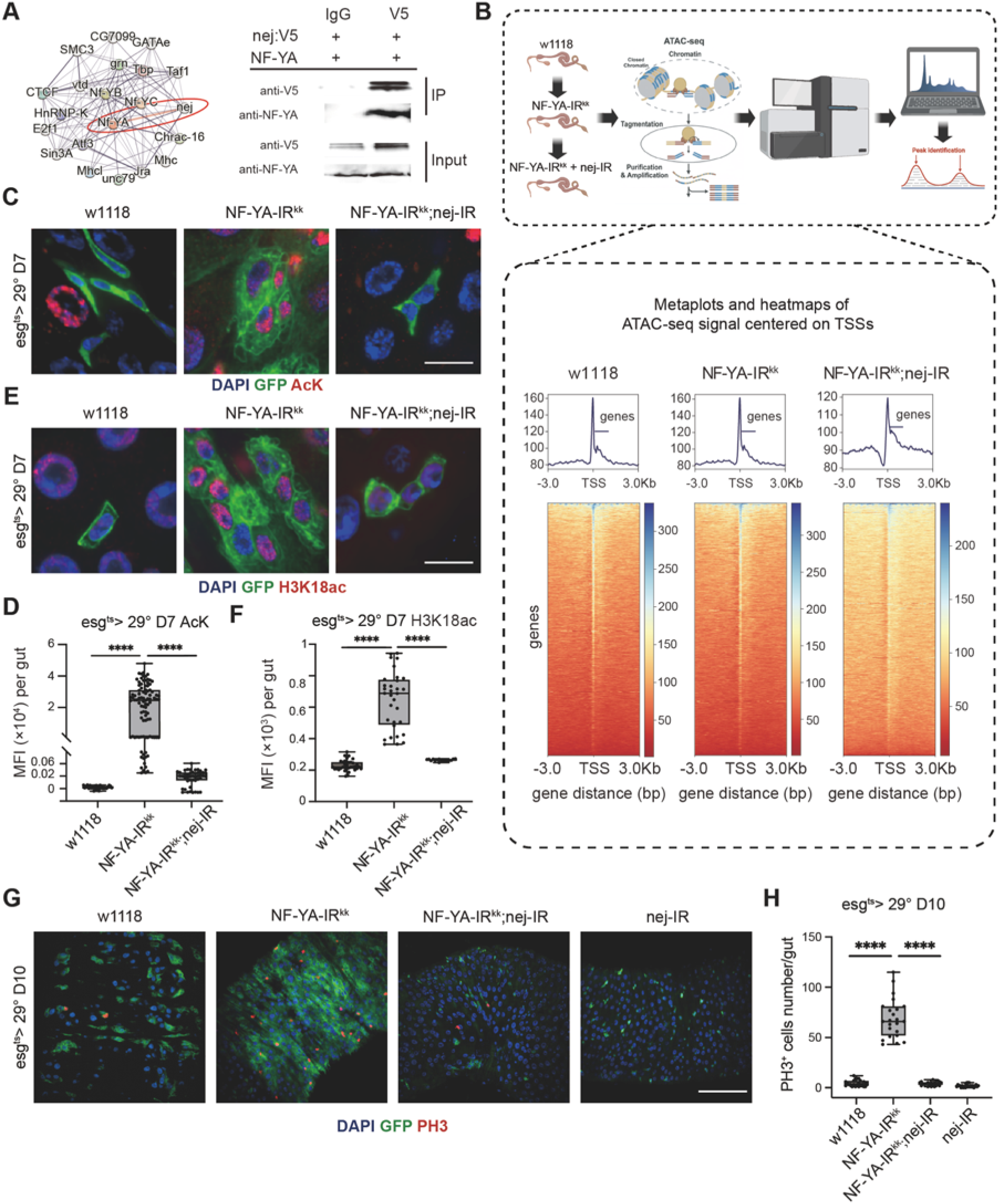
NF-YA depletion induces nej-mediated chromatin hyperacetylation and ISC hyperproliferation. (A) NF-YA physically interacts with the acetyltransferase Nej. (Left) STRING protein association network centered on NF-YA, highlighting functionally connected chromatin regulators including Nej. (Right) Co-immunoprecipitation (Co-IP) assay confirming the in vivo physical interaction between NF-YA and V5-tagged Nej. (B) Schematic view of the ATAC-seq experimental design (top) and metaplots/heatmaps (bottom) of ATAC-seq signals centered on transcription start sites (TSSs). NF-YA depletion (NF-YA-IR) increases promoter-proximal chromatin accessibility, whereas concurrent knockdown of nej (NF-YA-IR; nej-IR) largely attenuates this abnormal open chromatin state. (C and D) Representative images (C) and quantification (D) of pan-acetyl-lysine (AcK) staining in esg+ progenitor cells of the indicated genotypes. NF-YA depletion markedly increases global lysine acetylation, which is suppressed by nej codepletion. Blue, DAPI; green, GFP; red, AcK. Scale bar, 10 μm. n ≥ 38 midguts per group. (E and F) Representative images (E) and quantification (F) of H3K18ac staining in esg+ progenitor cells. Loss of NF-YA causes a strong increase in H3K18 acetylation, which is effectively reversed by nej RNAi. Blue, DAPI; green, GFP; red, H3K18ac. Scale bar, 10 μm. n ≥ 18 midguts per group. (G and H) Representative images (G) and quantification (H) of PH3+ mitotic cells in midguts from control (w1118), NF-YA-IR, and NF-YA-IR; nej-IR flies. nej depletion robustly suppresses the ISC hyperproliferation caused by NF-YA loss. Blue, DAPI; green, GFP; red, PH3. Scale bar, 50 μm. N ≥ 22 midguts per group. (Statistical note for D, F, H: Data are presented as box and whisker plots indicating the median, 25th – 75th percentiles, and minimum/maximum values. analyzed by one-way ANOVA with Dunnett’s multiple comparisons test; *P < 0.05,**P < 0.01, ***P < 0.001, ****P < 0.0001).

We next asked whether loss of NF-Y alters histone acetylation status in ISC, as nej/p300 is the main mediator for histone hyperacetylation to enable transcription activation (52). Our immunostaining with a pan acetyl lysine (AcK) or H3K18ac antibody revealed a marked increase in global lysine acetylation in NF-YA depleted ISCs relative to controls, and this elevation was strongly suppressed by nej co-depletion (Fig. 5 *C-F*). In contrast, knockdown of Nej alone caused a marked reduction in acetylation levels within stem cells. The efficiency of Nej depletion was further validated using two independent RNAi lines (Fig. S5 *C-F*). These data indicate that NF-Y loss induces a Nej dependent hyperacetylated chromatin state in ISCs.

We next examined whether this chromatin status functionally contributes to the excessive proliferation caused by NF-Y depletion. Interestingly, nej RNAi strongly suppressed NF-Y depletion induced ISC hyperproliferation, restoring mitotic activity toward control levels, whereas nej RNAi alone had little effect on basal proliferation (Fig. 5 *G* and *H*). These findings place Nej functionally downstream of, or in parallel with, NF-Y in controlling ISC proliferation. Collectively, these results identify Nej as a critical mediator of the chromatin state together with NF-Y to regulate transcription activation in ISC.

### NF-Y and nej coordinate the transcription activation of EGFR signaling to control stem cell activity

We next sought to identify the downstream effector pathway that contribute to the NF-Y/Nej mediated ISC overactivation. As our integrated RNA-seq and CUT&Tag analyses highlighted EGFR/MAPK associated genes as candidate direct NF-Y targets, we first examined chromatin accessibility across this pathway. ATAC-seq signal centered on transcription start sites (TSSs) of EGFR pathway genes revealed a pronounced increase in promoter-proximal accessibility upon NF-YA depletion, whereas co-depletion of nej markedly attenuated this increased genomic accessibility (Fig. 6 *A*). These findings indicate that NF-Y depletion induces EGFR/MAPK pathway transcription activation is dependent on nej modulated chromatin remodeling.

**Figure 6.**
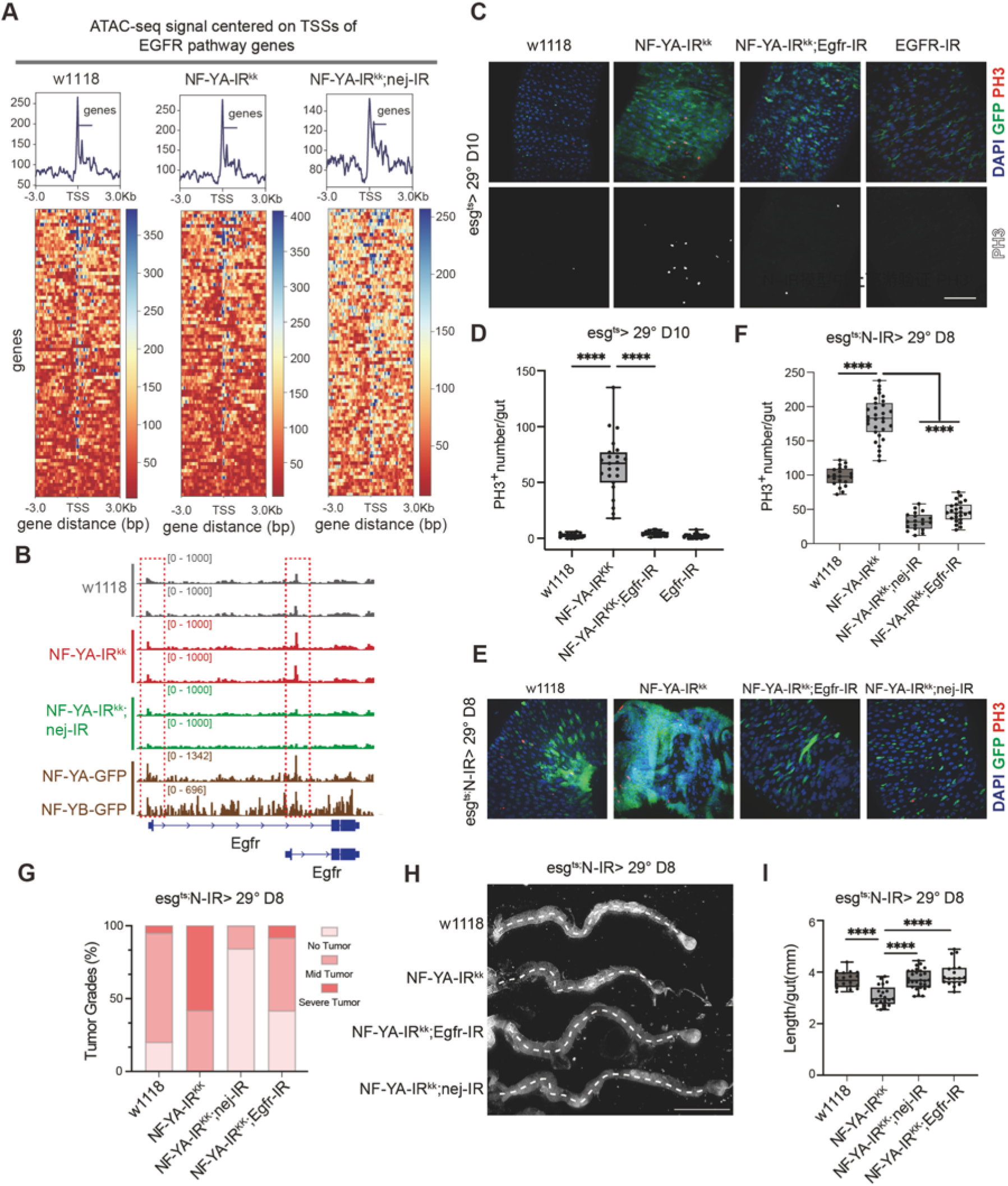
The NF-Y/Nej/EGFR axis controls intestinal stem cell activity and tumor progression.(A) Metaplots and heatmaps of ATAC-seq signal centered on transcription start sites (TSSs) of EGFR pathway genes in midguts from w1118, NF-YA-IR^kk^, and NF-YA-IR^kk^;nej-IR flies. NF-YA depletion increased promoter-proximal chromatin accessibility across EGFR pathway genes, whereas nej-RNAi attenuated this increased genomic accessibility. (B) Representative genome browser tracks at the Egfr locus showing ATAC-seq signals in w1118, NF-YA-IR^kk^, and NF-YA-IR^kk^;nej-IR midguts, together with NF-YA-GFP and NF-YB-GFP CUT&Tag tracks. Red dashed boxes highlight regions with differential accessibility and NF-Y binding. (C and D) Representative images (C) and quantification (D) of PH3+ mitotic cells in midguts from w1118, NF-YA-IR^kk^, NF-YA-IR^kk^;Egfr-IR, and Egfr-IR flies. Egfr-RNAi strongly suppressed the ISC proliferation induced by NF-YA depletion, whereas Egfr depletion alone had little effect on basal mitotic activity. Blue, DAPI; green, GFP; red, PH3. Scale bar, 50 μm. n ≥ 17 midguts per group. (E and F) Representative images (E) and quantification (F) of PH3+ mitotic cells in the Notch depletion (esg^ts^>N-IR) intestinal tumor model. NF-YA knockdown exacerbated tumor associated hyperproliferation, which was robustly rescued by codepletion of either Egfr or nej. n≥19 midguts per group, Blue, DAPI; green, GFP; red, PH3. Scale bar, 50 μm. (G) Quantification of tumor grades across the indicated genotypes in the esg^ts^>N-IR background. n≥19 midguts per group. (H and I) Representative whole-gut images (H) and quantification of midgut length (I) across the indicated genotypes in the esg^ts^>N-IR background. The shortened gut phenotype induced by tumor progression was further aggravated by NF-YA knockdown and restored by Egfr or nej codepletion. n≥19 midguts per group, Scale bar, 1 mm. (Statistical note for D, F, I: Data are presented as box and whisker plots indicating the median, 25th – 75th percentiles, and minimum/maximum values. analyzed by one-way ANOVA with Dunnett’s multiple comparisons test; *P < 0.05, **P < 0.01, ***P < 0.001, ****P < 0.0001)

A closer examination of the EGFR locus using integrated CUT&Tag and ATAC-seq analyses revealed that NF-YA depletion increased chromatin accessibility at regulatory regions of candidate genes (EGFR), whereas nej-RNAi attenuated this effect (Fig. 6 *B*). Notably, these accessible regions overlapped with NF-YA-GFP and NF-YB-GFP CUT&Tag peaks, supporting direct NF-Y complex occupancy at the Egfr locus. Similar patterns were observed at additional MAPK pathway genes, including Mkp3, Raf, Ras, and pnt, where NF-Y bound regions showed increased accessibility upon NF-YA knockdown and reduced accessibility after nej-RNAi (Fig. S6 *C*). These results indicate that multiple components of the EGFR/MAPK pathway are bona fide targets of the NF-Y/Nej regulatory module.

We next tested whether EGFR signaling is functionally required for the ISC hyperproliferation caused by NF-Y depletion. Coexpression of Egfr-RNAi with NF-YA-RNAi strongly suppressed ISC proliferation and restored mitotic activity toward control levels, whereas Egfr RNAi alone had little effect on basal proliferation (Fig. 6 *C* and *D*). These data identify EGFR/MAPK signaling as a key downstream effector of NF-Y loss induced ISC overproliferation.

Given that the NF-Y/Nej/EGFR axis restricts basal ISC proliferation, we asked whether it also influences intestinal tumorigenesis. Using a Notch depletion (*esg^ts^>N-IR*) induced tumor model, we found that knocking down NF-YA significantly exacerbated tumor progression, as evidenced by profoundly elevated ISC mitotic activity (Fig. 6 *E* and *F*), a shift toward severe tumor grades (Fig. 6 *G*), and a dramatic reduction in gut length typical of severe hyperplasia (Fig. 6 *H* and *I*). Importantly, co-depletion of either Egfr or nej in this background strongly suppressed these severe tumor phenotypes, restoring cellular proliferation and gut morphology to baseline tumor levels. These findings establish that the NF-Y/Nej/EGFR axis is not only critical for ISC homeostasis but also acts as a potent driver of tumor overgrowth.

We then asked whether NF-YA and Egfr expression are correlated across independent regenerative and pathological contexts. Spearman correlation analyses revealed a positive association between NF-YA and Egfr expression in the RNA-seq analysis including DSS induced regeneration, Ecc15 and Pseudomonas entomophila (Pe) infection, as well as the tumor conditions (Fig. S6 *A*). Consistent with this, correlation analyses of Mkp3, a negative-feedback regulator and canonical readout of EGFR/MAPK pathway activity(53), also supported coordinated regulation with NF-YA across these contexts (Fig. S6 *B*). These analyses suggest that expression of NF-YA is correlated with expression of the EGFR/MAPK genes across intestinal stress and tumor contexts

These findings hence position EGFR/MAPK signaling as a central downstream effector of NF-Y/nej mediated ISC quiescence. NF-Y depletion facilitates Nej dependent chromatin remodeling at EGFR/MAPK associated loci, thereby inducing proliferative genes expression and driving consequent ISC hyperproliferation. Together, our genetic, transcriptomic, chromatin accessibility and genomic loci enrichment analyses support a working model in which the NF-Y complex restrains Nej dependent accessibility on proliferative target genes to prevent ISC hyperactivation and preserve intestinal homeostasis, whereas loss of NF-Y releases this constraint and promotes dysplastic progression through activation of the EGFR/MAPK signaling (Fig. 7).

**Figure 7.**
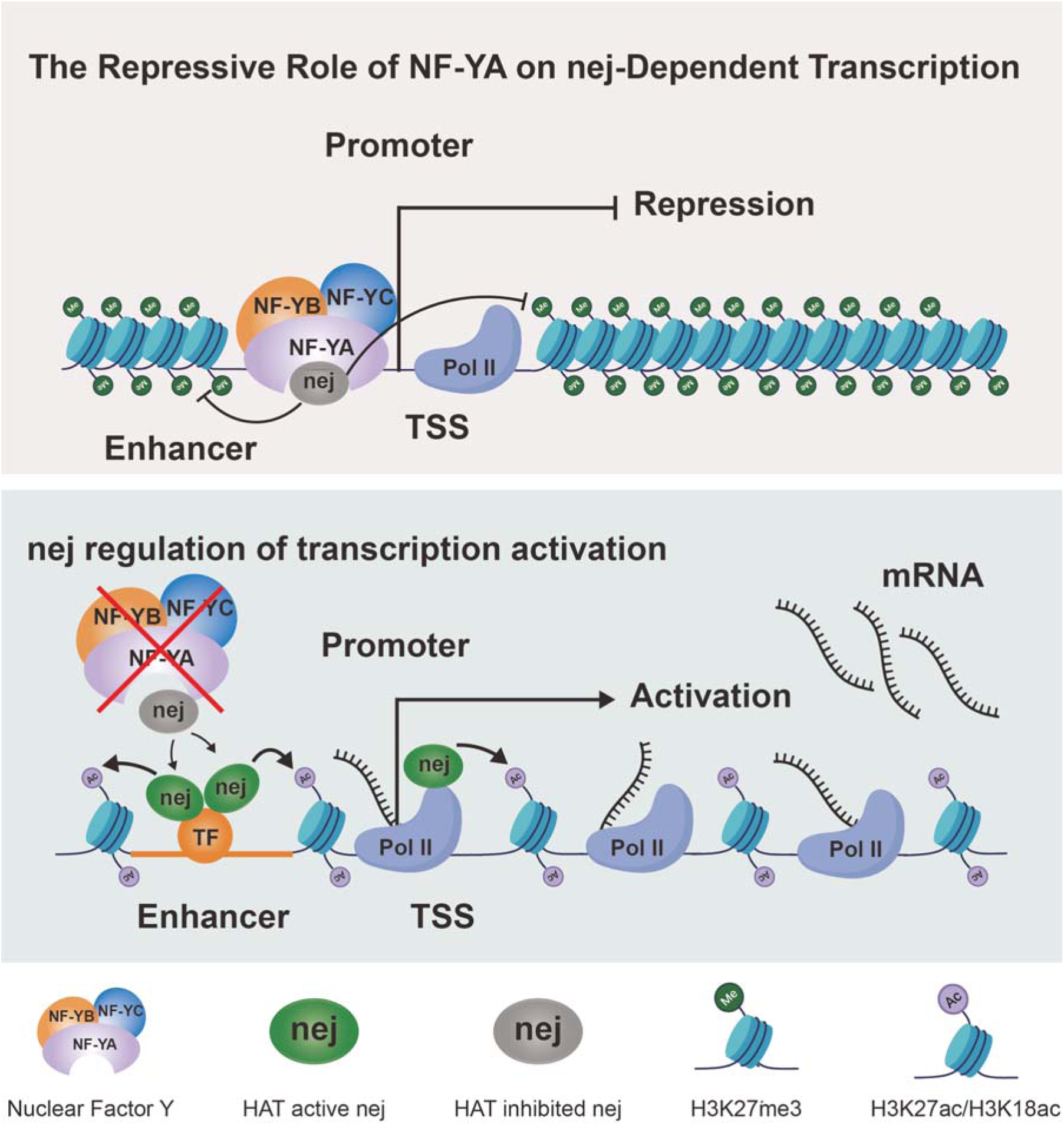
Proposed model for NF-Y-mediated repression of nej dependent transcriptional activation in ISC. Under homeostatic conditions, the NF-Y complex occupies promoter proximal regions of growth associated target genes and acts together with nej to restrain nej dependent chromatin activation, thereby limiting their transcriptional output. Upon NF-Y depletion, Nej promotes histone acetylation and chromatin opening at regulatory regions, leading to increased expression of proliferation promoting genes, including key components of the EGFR/MAPK pathway, and ultimately driving ISC hyperproliferation and dysplastic progression.

## Discussion

Precise control of adult stem cell activity is fundamental to tissue homeostasis, requiring a delicate balance between activation for regeneration and suppression to prevent hyperplasia. While the mechanisms that initiate stem cell proliferation following injury are well characterized including EGFR, JNK, JAK-STAT, and InR signaling pathways (10, 14, 15, 19), the negative feedback loops that terminate these responses and restore quiescence remain comparatively less understood. Our study identifies Nuclear Factor Y (NF-Y), in complex with the histone acetyltransferase Nejire (Nej)/p300, as a critical negative feedback module that restricts intestinal stem cell (ISC) proliferation by directly repressing expression of the EGFR signaling component EGFR, Mkp3, pointed. This work establishes a paradigm where active transcriptional repression is essential for re-establishing stem cell quiescence following regenerative episodes.

### Stem cell negative feedback regulation

The concept of active negative feedback is emerging as a conserved theme in stem cell biology across species. In the Drosophila midgut, several negative regulatory mechanisms have been identified that constrain ISC proliferation. The BMP pathway, acting as a niche derived signal, restrains ISC division to maintain homeostatic balance (4, 20, 54). Our findings add a new layer of complexity by demonstrating that NF-Y/Nej operates upstream of these effectors to transcriptionally gate the expression of EGFR, Mkp3, Raf, Ras, Pointed core components in the growth promoting cascade. This positions NF-Y as a key node that integrates transcriptional control with growth signaling, ensuring that the regenerative response is inherently self limiting. The requirement for robust negative feedback becomes particularly evident following tissue damage, when ISCs rapidly exit quiescence to replenish lost cells. During this regenerative phase, damaged enterocytes release mitogenic signals, including EGFR ligands, which drive ISC proliferation through Ras/MAPK cascades. Upon resolution of damage, the proliferative response must be terminated to prevent excessive growth. Failure to reestablish quiescence leads to persistent hyperplasia and can progress to dysplastic lesions and tumor like structures(11, 12, 22, 55). Our observation that NF-Y preferentially occupies the promoter of EGFR components to facilitate a repressive chromatin state via Nej/p300 suggests a highly targeted mechanism for resetting the system after regeneration. Dysregulation of negative feedback circuits has profound implications for disease. The loss of NF-Y in our model enhances ISC proliferation and exacerbates stress and tumor induced mortality, mirroring the consequences of chronic, uncontrolled stem cell activity seen in conditions like inflammatory bowel disease and colorectal cancer. In these human pathologies, sustained activation of pathways such as EGFR, InR, and JAK-STAT disrupts homeostatic balance and promotes tumorigenesis (16, 25). The evolutionary conservation of NF-Y, p300, and the EGFR axis from flies to mammals suggests that this regulatory mechanism is ancient and potentially targetable. By elucidating how NF-Y/Nej orchestrates the termination of growth signaling, our study provides a mechanistic framework for understanding how stem cell quiescence is reestablished. Future investigations in mammals should be explored about how this transcriptional repressive complex is itself regulated by upstream signals from the niche and how its dysfunction contributes to age related decline in regenerative capacity and cancer initiation.

## Materials and Methods

### Drosophila melanogaster strains and culture

All fly stocks were maintained on standard cornmeal yeast agar medium at 25°C unless otherwise specified. For temporal control of transgene expression in intestinal progenitor cells, the esg-Gal4, UAS-GFP, tub-Gal80^ts^ system (hereafter referred to as esg^ts^) was used. Crosses were maintained at 18°C to suppress Gal4 activity during development. For enterocyte specific manipulations, the Myo1A-Gal4, UAS-GFP, tub-Gal80ts system (hereafter referred to as Myo1A^ts^) was used. Following eclosion, 3 day old adult progeny were shifted to 29°C to induce expression of UAS driven transgenes.For tumor model experiments, progenitor specific tumors were induced using the esg^ts^ system. The Notch depletion tumor model was generated by expressing UAS-Notch-RNAi in esg+ intestinal progenitor cells. The Ras/Apc tumor model was generated by co expressing UAS-Ras^V12^ and UAS-Apc-RNAi in the same compartment.The following fly lines were used in this study: w1118 (control); esg-Gal4, UAS-GFP, tub-Gal80^ts^; Myo1A-Gal4, UAS-GFP, tub-Gal80^ts^; UAS-mCherry; Delta-LacZ; endogenous NF-YA-GFP, NF-YB-GFP, and NF-YC-GFP lines; UAS-NF-YA-RNAi^kk^, UAS-NF-YB-RNAi^kk^, UAS-NF-YB-RNAi^trip^, UAS-NF-YC-RNAi^trip^, UAS-nej-RNAi^kk^, UAS-Egfr-RNAi^trip^, UAS-Notch-RNAi, UAS-Ras^V12^, UAS-Apc-RNAi, UAS-Cas9, sepia-gRNA gRNA-NF-YA, gRNA-NF-YC, and the esg ts flip/out lineage tracing stock. Fly stocks were obtained from the Bloomington Drosophila Stock Center (BDSC), the Vienna Drosophila Resource Center (VDRC), the Tsinghua Fly Center, or were generous gifts from other laboratories, as indicated in the Key Resources Table.

### DSS-induced intestinal injury time-course for PH3 analysis

For the intestinal injury time course experiment shown in Fig. 1A, 3 to 5 day old adult female flies were fed 3% dextran sulfate sodium (DSS; MP Biomedicals, 02160110-CF) dissolved in 5% sucrose using a filter paper feeding assay. Briefly, 200 μL of DSS solution was applied to a filter paper placed on the surface of the food vial and replaced daily. Flies were maintained on DSS for 5 days and then transferred back to standard cornmeal yeast agar food for recovery. Midguts were dissected at the indicated time points from day 1 to day 8, and mitotic activity was assessed by phospho-histone H3 (PH3) staining.

### DSS treatment and recovery paradigm for transcriptomic profiling

For the bulk RNA-seq experiment shown in Fig. 1B, 3 to 5 day old adult female flies were fed 3% DSS dissolved in 5% sucrose using the same filter paper feeding method for 4 days. Midguts were collected at DSS day 1 (D1) and DSS day 4 (D4) during the injury phase, and at recovery day 1 (corresponding to D5) and recovery day 4 (corresponding to D8) after transfer to standard food. Control samples were collected from flies fed 5% sucrose alone at DSS D1 and DSS D4 using the same feeding scheme. Dissected midguts were used for downstream transcriptomic analysis.

### DSS challenge in genetically manipulated flies

For DSS challenge experiments in transgenic flies, adult females were first maintained at 29°C for 7 days to induce transgene expression and were then transferred to 3% DSS dissolved in 5% sucrose for an additional 3 days prior to dissection or survival analysis.

### Immunofluorescence staining and confocal microscopy

Adult female midguts were dissected in 1× phosphate buffered saline (PBS; Solarbio) and fixed in 4% paraformaldehyde (PFA) for 25 min at room temperature. Tissues were washed four times in 1× PBST (1× PBS containing 0.1% Triton X-100; Solarbio) and blocked for 30 min in blocking solution (1% bovine serum albumin [BSA; Solarbio] in 1× PBST). Midguts were then incubated with primary antibodies diluted in blocking solution at 4°C overnight, followed by incubation with the appropriate secondary antibodies for 2 hr at room temperature. Samples were subsequently washed four times in 1× PBST and mounted in Vectashield mounting medium containing DAPI (Vector Labs).

Primary antibodies used in this study included rabbit anti-PH3 (1:800, Cell Signaling Technology, 9701L), rabbit anti-AcK (1:3000, Cell Signaling Technology, 9814S), and rabbit anti-H3K18ac (1:3000, Active Motif, 39755). Secondary antibodies used were goat anti-rabbit Alexa Fluor 488 (1:3000, Thermo Fisher Scientific, A11034) and donkey anti-rabbit Alexa Fluor 555 (1:3000, Thermo Fisher Scientific, A31572), as indicated for individual experiments. Images were acquired using a Zeiss LSM 980 confocal microscope under identical acquisition settings across experimental groups and processed using ImageJ/Fiji software. For comparative imaging experiments, control and experimental samples were processed in parallel and imaged using identical laser power, gain, and exposure settings.

### Survival analysis

For survival assays, 100–200 female flies per genotype were collected within 24 hr of eclosion and maintained at 29°C under the indicated experimental conditions. Flies were housed in standard food vials and transferred to fresh food or treatment vials every 2 days unless otherwise specified. Dead flies were scored daily until all flies had died.

For survival experiments performed under intestinal injury conditions, flies were first maintained at 29°C for transgene induction as indicated and were then transferred to 3% DSS for the duration specified in the corresponding experiment. For tumor model survival assays, flies carrying the indicated transgenes were maintained continuously at 29°C after adult induction.

Survival curves were plotted using the Kaplan–Meier method, and statistical significance between groups was determined using the log-rank (Mantel–Cox) test.

### Smurf assay

To assess intestinal barrier integrity, flies were transferred to food containing 2% Acid Blue 9 (Shanghai Dibai, MK30) for 24 hr. Flies showing blue dye leakage outside the intestinal tract were scored as Smurf positive.

### Gut morphology and length measurement

For whole gut morphology analysis, dissected intestines were imaged under a stereomicroscope or low magnification brightfield microscope. Gut length was measured in ImageJ/Fiji along the central longitudinal axis using the segmented line tool. Where indicated, gut length was normalized to body mass as specified in the corresponding figure legends.

### CRISPR/Cas9 mediated gene knockout

For temporal knockout experiments, esg^ts^ > Cas9 flies carrying sepia-gRNA, gRNA-NF-YA, or gRNA-NF-YC were shifted to 29°C for 5 days to induce mutagenesis in intestinal progenitor cells. Flies were then returned to the permissive temperature (18°C) for 30 days to allow epithelial turnover before analysis.

### Protein extraction, Co-Immunoprecipitation (Co-IP), and Western blotting

For in vivo protein extraction, adult Drosophila midguts or whole flies of the indicated genotypes were dissected or collected in ice cold PBS, respectively. Tissues were homogenized in standard RIPA lysis buffer (50 mM Tris-HCl pH 7.4, 150 mM NaCl, 1% NP-40, 0.5% sodium deoxycholate, 0.1% SDS) supplemented with a complete protease and phosphatase inhibitor cocktail (Roche). Lysates were incubated on ice for 30 min and centrifuged at 12,000 × g for 15 min at 4°C to remove cellular debris.

For Co-IP assays, pre-cleared equal amounts of total protein lysates were incubated with specific primary antibodies overnight at 4°C with gentle rotation. Protein A/G magnetic beads (36427ES03) were added and incubated for an additional 2 hr at 4°C. The beads were then washed five times with ice cold lysis buffer to remove no specifically bound proteins. Bound protein complexes were eluted by boiling in 1× SDS loading buffer at 95°C for 10 min.

For Western blotting, equal amounts of input lysates and immunoprecipitated samples were resolved by SDS-PAGE and transferred onto PVDF membranes (Millipore). Membranes were blocked with 5% non-fat milk in TBST (Tris buffered saline containing 0.1% Tween-20) for 1 hr at room temperature, followed by overnight incubation at 4°C with the following primary antibodies: anti-V5 (1:200, F2690) and anti-NF-YA (1:200, 12981-1-AP). After three washes with TBST, membranes were incubated with HRP conjugated secondary antibodies (1:5000) for 1 hr at room temperature. Protein bands were visualized using an Enhanced Chemiluminescence (ECL) and captured using a digital imaging system.

### RNA-seq and bioinformatic analysis

For bulk RNA-seq, 10 dissected adult female midguts were collected per sample, with three independent biological replicates for each condition. Freshly dissected midguts were kept on dry ice, and shipped to a commercial sequencing provider for RNA extraction, library construction, and sequencing.

Raw sequencing data were analyzed in house. Paired end reads were mapped to the Drosophila melanogaster dm6 reference genome from FlyBase using HISAT2 (version 2.2.1). Read counting was performed using FeatureCounts (version 2.0.6). Differentially expressed genes (DEGs) were identified using the R package DESeq2. Unless otherwise specified, genes with log2 fold change > 1 and adjusted pvalue < 0.05 were considered significantly differentially expressed.

Gene Ontology (GO) and KEGG pathway enrichment analyses were performed using clusterProfiler. Gene Set Enrichment Analysis (GSEA) was conducted using GSEA software (version 4.3.3). Principal component analysis (PCA), temporal clustering, overlap analysis, and heatmap visualization were performed in R. Clustering visualization was generated using Cluster GVis where applicable.

For analysis of publicly available RNA-seq datasets, raw or processed data were obtained from the corresponding repositories as indicated in the manuscript. When raw sequencing files were available, paired end reads were processed using the same pipeline described above, including alignment to dm6 with HISAT2, quantification with Feature Counts, and differential expression analysis with DESeq2. Correlation analyses and additional visualization were performed in R.

### CUT&Tag assay

CUT&Tag was performed using the CUT&Tag reagent kit (Ruoyu Biotech, CUT-01) according to the manufacturer’s instructions. 30 midguts were dissected in ice-cold sterile PBS. Gut tissues were cut into 3-4 pieces and transferred into low adhesion microcentrifuge tubes, followed by gentle washing with 400 μL cold PBS to remove debris.

For tissue dissociation, gut fragments were incubated in 500 μL of 0.5% trypsin (Solarbio, T1302) for 15 min at 29°C with shaking at 1000 rpm. After brief settling on ice, the supernatant was collected and centrifuged at 500 × g for 5 min, and the pellet was resuspended in 50 μL cold PBS. This digestion step was repeated 2-3 times until complete dissociation was achieved. The pooled cell suspensions were filtered through 100 μm cell strainers and counted using a hemocytometer with 0.4% Trypan blue staining.

Next, 5 μL of preactivated ConcanavalinA coated beads were added to the cell suspension. Cells were resuspended in 100 μL Dig wash Buffer and incubated sequentially with primary and secondary antibodies. Primary antibodies used were anti-GFP (1:100, Proteintech, 50430-2-AP) and anti-H3K4me3 (1:100, Cell Signaling Technology, 9751). A 1:250 dilution of the pAG-Tn5 adapter complex was prepared in Dig-wash Buffer and added to the samples. After resuspension in Fragmentation Buffer containing MgCl₂, the fragmentation reaction was terminated by addition of 2.5 μL EDTA to the 50 μL bead mixture. DNA was purified using magnetic beads, PCR amplified, and subjected to library construction. Final libraries were sequenced on an Illumina NovaSeq 150PE platform.

### CUT&Tag data analysis

Raw sequencing reads were preprocessed using Trimmomatic (version 0.39) to remove low quality bases and adapter sequences. Filtered reads were aligned to the Drosophila melanogaster dm6 reference genome using Bowtie2. Peak calling was performed with MACS2 (version 2.2.9.1) using a qvalue cutoff < 0.01. Peak annotation was carried out using the R package ChIPseeker.

Functional enrichment analysis of CUT&Tag target genes was performed using PANGEA. For visualization of genomic occupancy around transcription start sites (±3 kb around TSSs), BAM files were converted to bigWig format using bamCoverage in deepTools with the normalize Using RPKM option, followed by generation of signal matrices and heatmaps using computeMatrix and plotHeatmap. Representative loci were visualized using a genome browser such as IGV.

### ATAC-seq assay

ATAC-seq was performed using the ATAC-Seq Library Preparation Kit (Ruoyu Biotech, ATS-01) according to the manufacturer’s instructions. Adult female flies of the indicated genotypes were collected at the appropriate time point, and dissected midguts were processed immediately in ice cold PBS. Intestinal tissues were dissociated into single cell suspensions by enzymatic digestion, and the resulting cells were washed and resuspended in the buffer system provided by the kit. Cell suspensions were then bound to ConcanavalinA coated beads and permeabilized in Dig buffer containing digitonin and BSA before transposition.

For the transposition reaction, bead bound cells were incubated with Tn5 transposase in TD buffer at 37°C for 30 min. Reactions were terminated with Stop buffer, and transposed DNA was purified using magnetic beads according to the kit protocol. Library pre-amplification was first performed to estimate the optimal number of PCR cycles, followed by final library amplification using indexed primers. PCR products were purified using 1.2× DNA purification beads, and the final libraries were quantified and assessed for fragment size distribution prior to sequencing. Libraries were sequenced on an Illumina NovaSeq 150PE platform.

### ATAC-seq data analysis

Raw ATAC-seq reads were processed to remove adapter sequences and low quality bases using Trimmomatic (version 0.39). Filtered reads were aligned to the Drosophila melanogaster dm6 reference genome using Bowtie2. Uniquely mapped reads were retained for downstream analysis.

Accessible chromatin peaks were identified using MACS2. Peak annotation was performed using the R package ChIPseeker. For genome wide accessibility profiling around transcription start sites (TSSs), BAM files were converted to bigWig format using the bamCoverage function in deepTools with RPKM normalization, followed by matrix generation and visualization using computeMatrix, plotHeatmap, and plotProfile. Genomic feature annotation of ATAC-seq peaks was used to determine promoter enriched accessibility patterns.

For pathway focused analysis, ATAC-seq signal was examined at EGFR/MAPK associated genes and other selected loci. Representative browser tracks were visualized using IGV or an equivalent genome browser.

### QUANTIFICATION AND STATISTICAL ANALYSIS

All statistical analyses were performed using GraphPad Prism and R unless otherwise specified. For comparisons between two groups, unpaired two tailed Student’s t tests were used. For comparisons among three or more groups, one-way ANOVA followed by Dunnett’s multiple-comparisons test was applied. Kaplan–Meier survival curves were analyzed using the log-rank (Mantel–Cox) test. Spearman correlation analysis was used for expression correlation analyses as indicated.Data are presented as mean ± SD for bar graphs, or as box-and-whisker plots (indicating the median, 25th–75th percentiles, and minimum/maximum values) as specified in the relevant figure legends. Unless otherwise stated, all experiments were performed with at least three independent biological replicates.Statistical significance was defined as follows:P < 0.05 (*),P < 0.01 (**),P < 0.001 (***), andP < 0.0001 (****).

## Data availability

CUT&Tag and ATAC data generated in this study are submitted to zenodo database 10.5281/zenodo.19478962. The RNA-seq data has been submitted to zenodo database 10.5281/zenodo.19477648.

If your research involved human or animal participants, please identify the institutional review board and/or licensing committee that approved the experiments. Please also include a brief description of your informed consent procure if your experiments involved human participants.

## Acknowledgments

We would like to thank B. Edgar, Y.Qi, XH. Lin, ZZ. Zhai, Developmental Studies Hybridoma Bank (DSHB), Bloomington Drosophila Stock Center, Kyoto stock center and Vienna Drosophila RNAi Center (VDRC) for fly stocks and antibodies. We thank Yansong Xiong, Yanan Hao and the Analytical Instrumentation Center of Hunan University for assistance in confocal microscopy. The work from Zhou Laboratory is supported by National Natural Science Foundation of China (32270890), the Department of Science and Technology of Hunan Province (x), and Hunan Provincial Key Laboratory of Anti-Resistance Microbial Drugs, the third hospital of Changsha (No:2023TP1013-08).

## Author Contributions

Yang SC, Luo CS, Peng GF conduct the experiments and interpret the results, write the manuscript. Zheng KW, Han K, Zhou J supervise the project, edit the manuscript and acquire funding.

## Supporting Information

**Figure S1.**
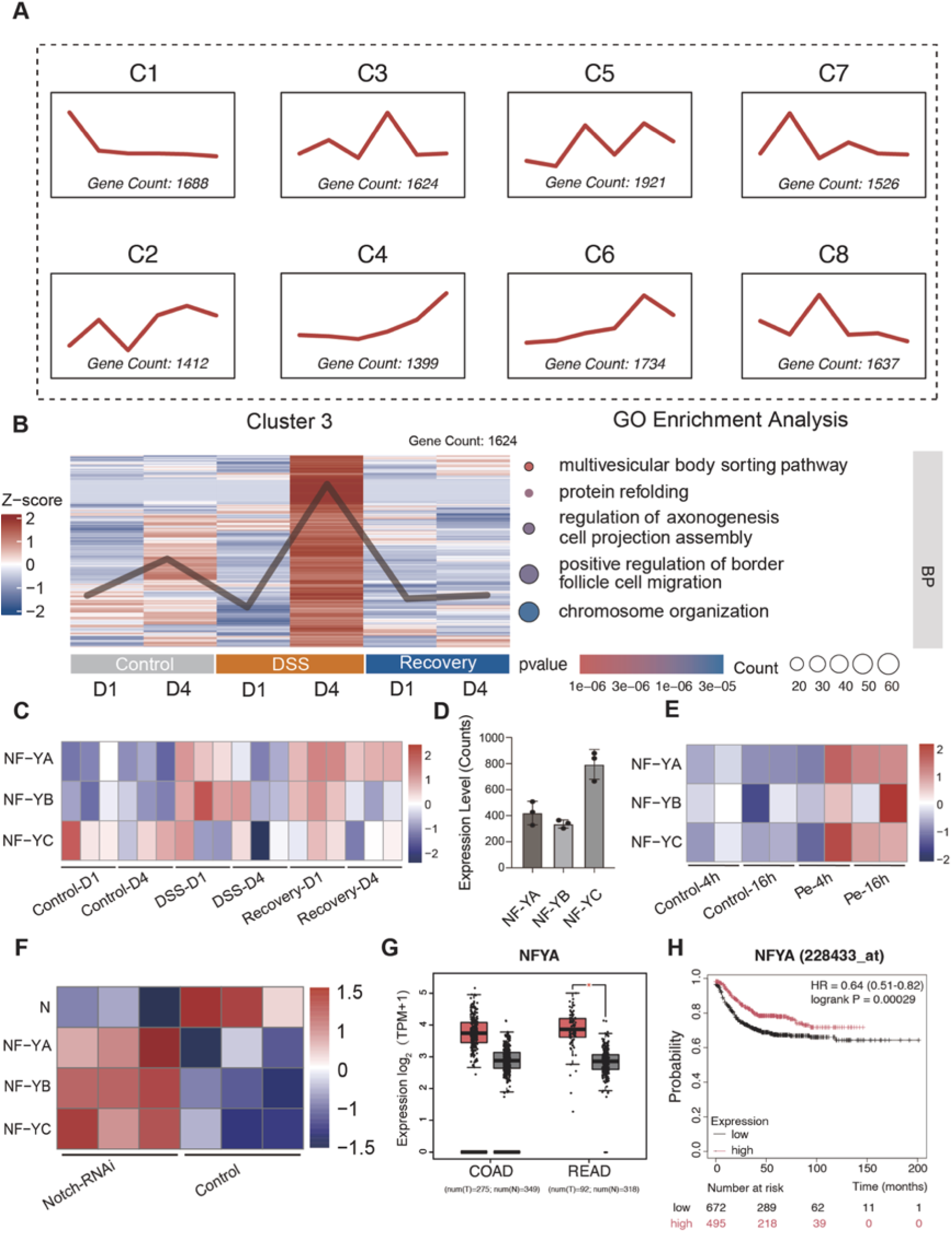
The NF-Y complex responds to multiple intestinal stresses and correlates with colorectal cancer prognosis. (A) Temporal coexpression clustering of regeneration-associated genes. Trend plots of eight gene clusters (C1–C8) defined by distinct expression dynamics during DSS injury and recovery. Clusters C3 and C5 display injury or recovery enriched patterns. Gene number per cluster is indicated. (B) Functional characterization of Cluster 3 genes. (Left) Heatmap and mean Z-score trend profile of Cluster 3 genes (n = 1,624). (Right) GO enrichment analysis of Cluster 3 genes; highlighted biological processes include multivesicular body sorting, protein refolding, regulation of axonogenesis, cell projection assembly, border follicle cell migration, and chromosome organization. Dot color indicates p value; dot size indicates gene count. (C) Heatmap showing normalized expression (Z-score) of NF-YA, NF-YB, and NF-YC across control (D1, D4), DSS treated (D1, D4), and recovery (Recovery-D1, Recovery-D4) stages. All three subunits are coordinately upregulated during DSS injury and remain elevated into early recovery. (D) Basal transcript levels of the NF-Y complex subunits. Bar graph showing the expression levels (RNA-seq counts) of NF-YA, NF-YB, and NF-YC in the adult midguts of control (w1118) flies. Data are represented as mean ± SD. Each dot represents an independent biological replicate (n = 3). (E) Induction of NF-Y subunits following enteric bacterial infection. Heatmap showing expression of NF-YA, NF-YB, and NF-YC at 4 h and 16 h after oral P. entomophila infection. (F) NF-Y upregulation in a Notch loss of function tumor model. Heatmap showing increased expression of NF-YA, NF-YB, and NF-YC in midguts with Notch depletion (Notch RNAi) compared with control. (G) Expression of NFYA subunits in human colorectal cancer. Box plots generated using GEPIA2 (TCGA and GTEx datasets) showing transcript levels (log₂(TPM+1)) in tumor (T; red) versus normal adjacent tissue (N; gray) for colon adenocarcinoma (COAD; T: n = 275, N: n = 349) and rectal adenocarcinoma (READ; T: n = 92, N: n = 318) from TCGA and GTEx datasets. *p < 0.05. (H) Kaplan–Meier analysis of overall survival in colorectal cancer patients stratified by NFYA mRNA expression level (probe 228433_at), generated using Kaplan–Meier Plotter. High NFYA expression (pink) is associated with significantly better overall survival compared with low expression (black). HR = 0.64 (0.51–0.82); log-rank p = 0.00029.

**Figure S2.**
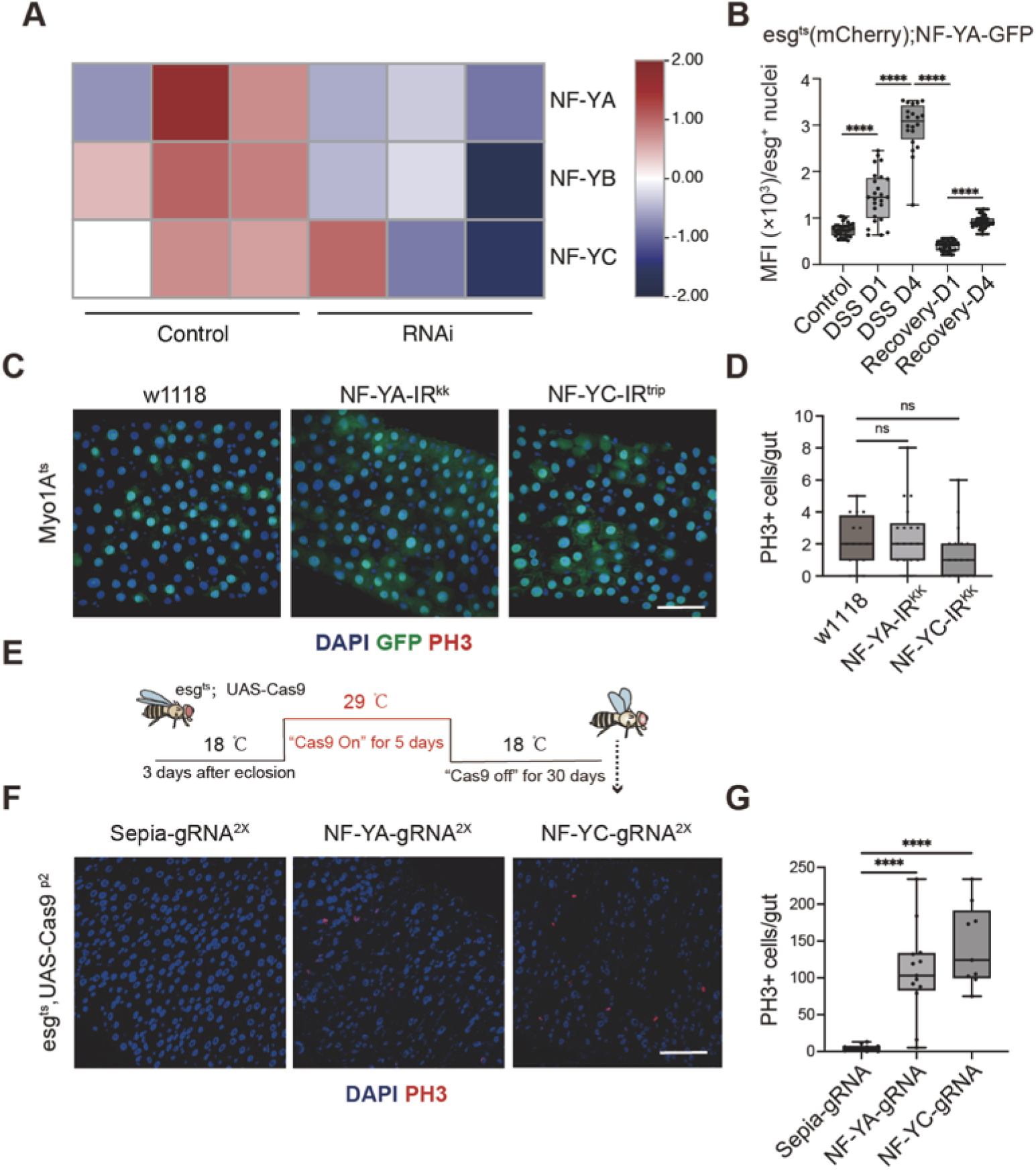
The NF-Y complex is required to maintain intestinal homeostasis. (A) Validation of RNAi knockdown efficiency. Heatmap showing the relative transcript levels of NF-YA, NF-YB, and NF-YC in midguts from control (w1118) and indicated RNAi lines. (B) Quantification of nuclear NF-YA-GFP mean fluorescence intensity (MFI) within esg+ nuclei across injury and recovery stages, complementing the total progenitor cellular signal shown in Figure 2B. ****p < 0.0001; analyzed by one-way ANOVA with Sidak’s multiple comparisons test (n ≥ 20 guts per time point). (C and D) EC specific depletion of NF-Y does not affect ISC proliferation. (C) Representative images of midguts expressing NF-YA or NF-YC RNAi driven by the Myo1Ats EC specific driver. PH3 (red) marks mitotic cells; GFP (green) marks ECs. Scale bar, 50 μm. (D) Quantification of PH3+ cells per gut. ns, not significant (p > 0.05); n ≥ 16 guts per group. (E–G) CRISPR/Cas9 mediated knockout of NF-Y subunits in ISCs reveals their requirement for maintaining intestinal homeostasis. (E) Schematic view of the temporal CRISPR/Cas9 knockout experimental design using esg^ts^ > UAS-Cas9. Cas9 expression was induced for 5 days post eclosion, followed by a 30 day recovery period before analysis. (F) Representative images of midguts stained for PH3 (red) and DAPI (blue) at 30 days after Cas9 induction in flies carrying gRNAs targeting sepia (control), NF-YA, or NF-YC. Scale bar, 50 μm. (G) Quantification of PH3+ cells per gut, demonstrating sustained ISC hyperproliferation following NF-Y ablation. ****p < 0.0001; n ≥ 9 guts per group. (Statistical note for D, G: Data are presented as box and whisker plots indicating the median, 25th–75th percentiles, and minimum/maximum values. analyzed by one-way ANOVA with Dunnett’s multiple comparisons test; *P < 0.05,**P < 0.01, ***P < 0.001, ****P < 0.0001)

**Figure S3.**
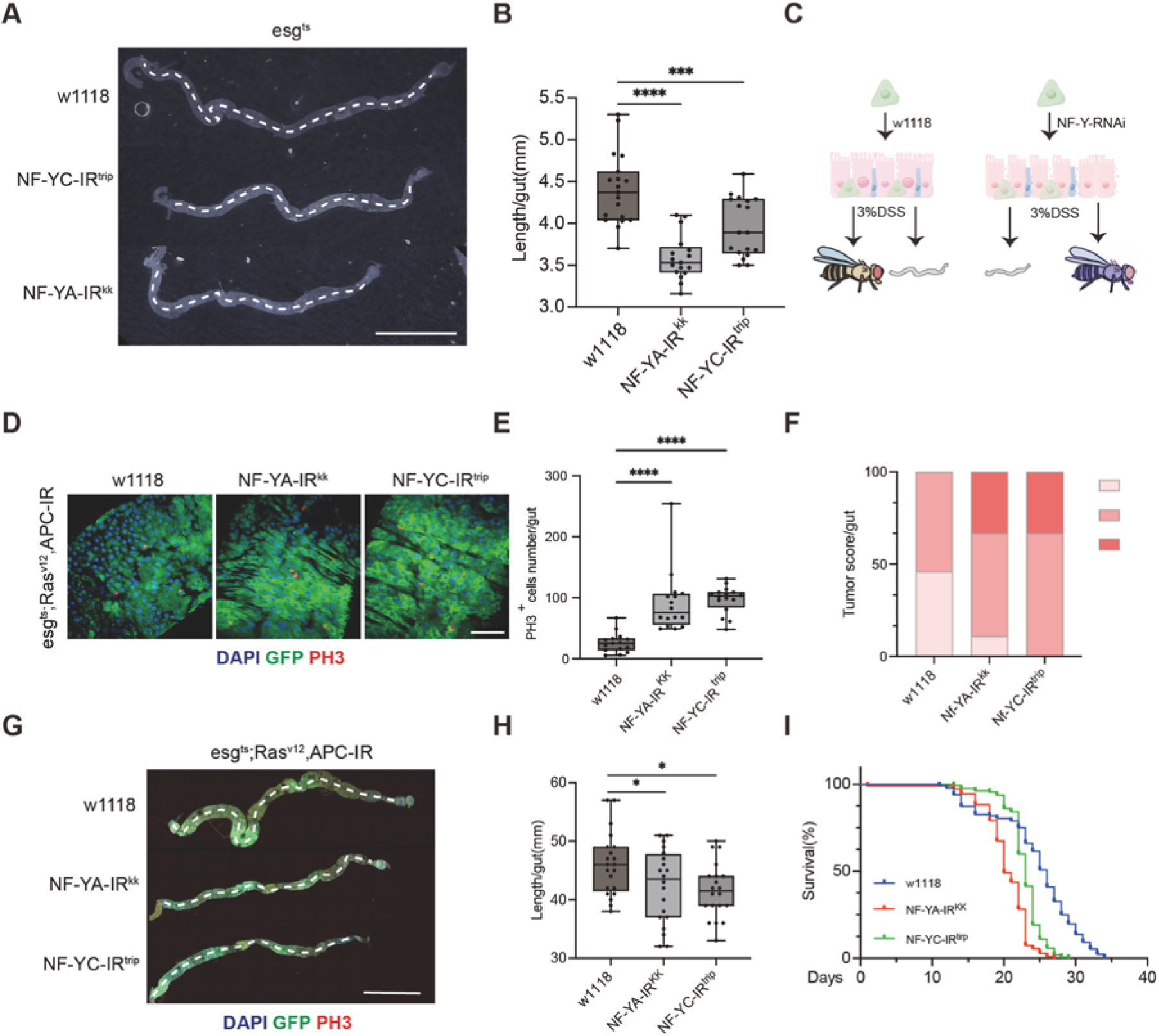
NF-Y depletion enhances Ras^V12^, Apc-IR-driven tumor progression. (A and B) Representative whole gut images (A) and gut length quantification (B) in control and NF-Y RNAi flies. NF-Y loss caused significant intestinal shortening. Dashed lines indicate measured gut length. n ≥17 midguts per group. (C) Schematic summary illustrating the increased susceptibility of NF-Y-RNAi intestines to 3% DSS challenge compared with control intestines. (D and E) Representative images (D) and quantification of PH3+ mitotic cells (E) in midguts expressing together with control, NF-YA RNAi, or NF-YC RNAi. NF-Y depletion further increased mitotic activity in the Ras^V12^, Apc-IR tumor model. Blue, DAPI; green, GFP; red, PH3. Scale bar = 50 µm; n ≥ 15 midguts per group. (F) Quantification of tumor-grade distribution in the Ras^V12^, Apc-IR background. NF-Y depletion shifted tumor burden toward more advanced grades. n ≥26 midguts per group. (G and H) Representative whole gut images (G) and gut length quantification (H) comparing Ras^V12^, Apc-IR flies with or without NF-Y RNAi. NF-Y loss further exacerbated gut shortening and structural distortion. Scale bar = 1 mm; n ≥20 midguts per group. (I) Kaplan-Meier survival analysis of flies of indicated genotypes. NF-Y knockdown significantly reduced survival of tumor flies. n ≥ 110 flies per genotype; log-rank test; ****P < 0.0001(Statistical note for B, E, H: Data are presented as box and whisker plots indicating the median, 25th–75th percentiles, and minimum/maximum values. analyzed by one-way ANOVA with Dunnett’s multiple comparisons test; *P < 0.05,**P < 0.01, ***P < 0.001,****P < 0.0001

**Figure S4.**
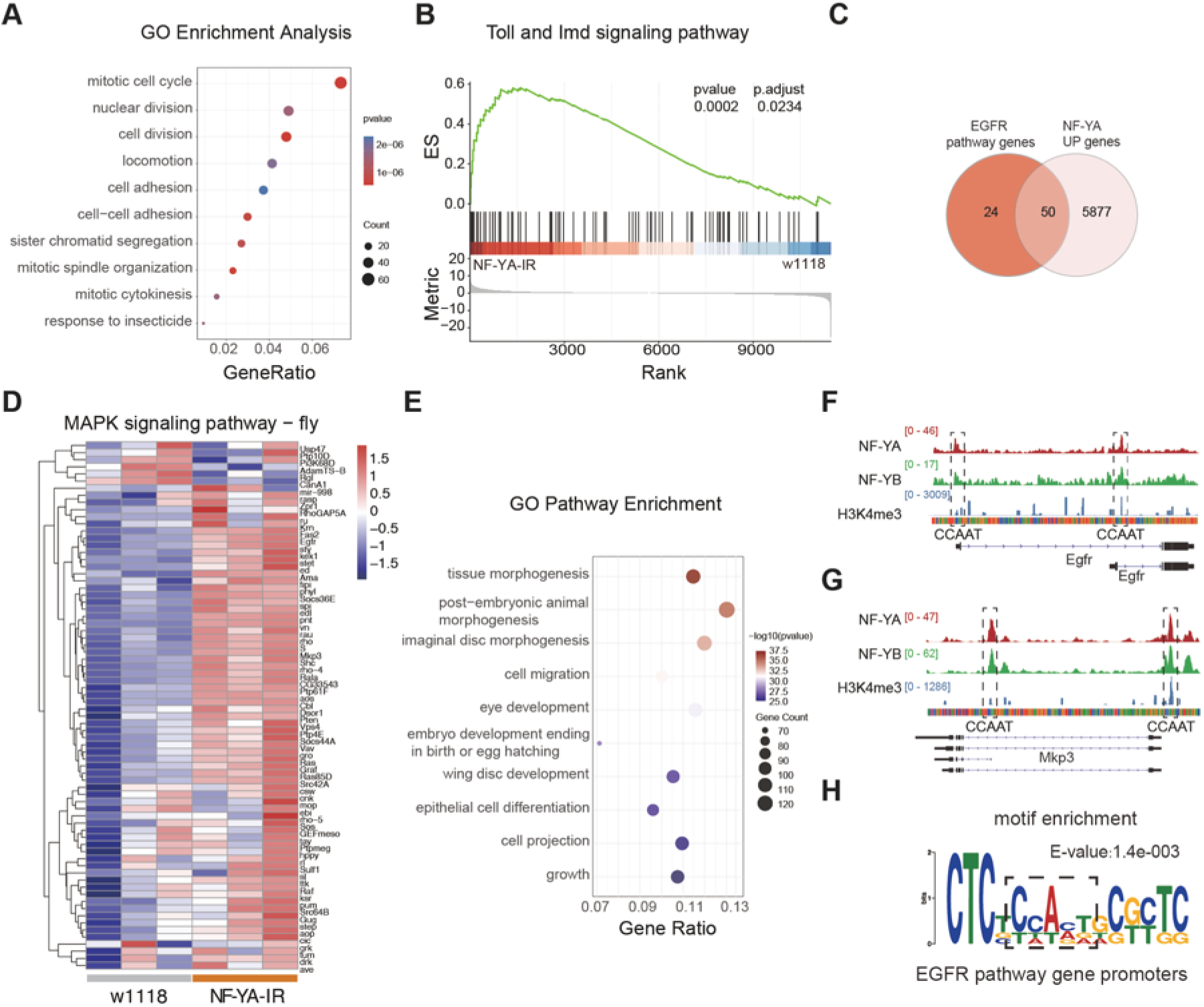
NF-Y targets transcriptional programs associated with mitosis, immunity, and EGFR/MAPK signaling. (A) Gene Ontology (GO) enrichment analysis of all DEGs identified between NF-YA depleted and control (w1118) midguts, highlighting processes related to mitosis and cell division. (B) GSEA plot showing significant positive enrichment of the Toll and Imd innate immune signaling pathway upon NF-YA depletion. (C) Venn diagram showing the overlap between genes upregulated by NF-YA depletion and genes curated in the EGFR pathway. (D) Heatmap showing the normalized expression (Z-score) of genes associated with the MAPK signaling pathway in control and NF-YA depleted midguts. (E) GO enrichment analysis of the 1,186 direct NF-Y target genes (defined in Figure 4I), showing enrichment for tissue morphogenesis and cell differentiation processes. (F and G) Representative genomic tracks of NF-YA, NF-YB, and H3K4me3 CUT&Tag signals at the loci of key MAPK pathway components Egfr (F) and Mkp3 (G). Dashed boxes indicate the precise colocalization of NF-Y peaks with predicted CCAAT box motifs at the promoter regions. (H) Motif enrichment analysis specifically within the promoters of EGFR pathway genes targeted by NF-Y, displaying the consensus binding sequence (E-value = 1.4e-03).

**Figure S5.**
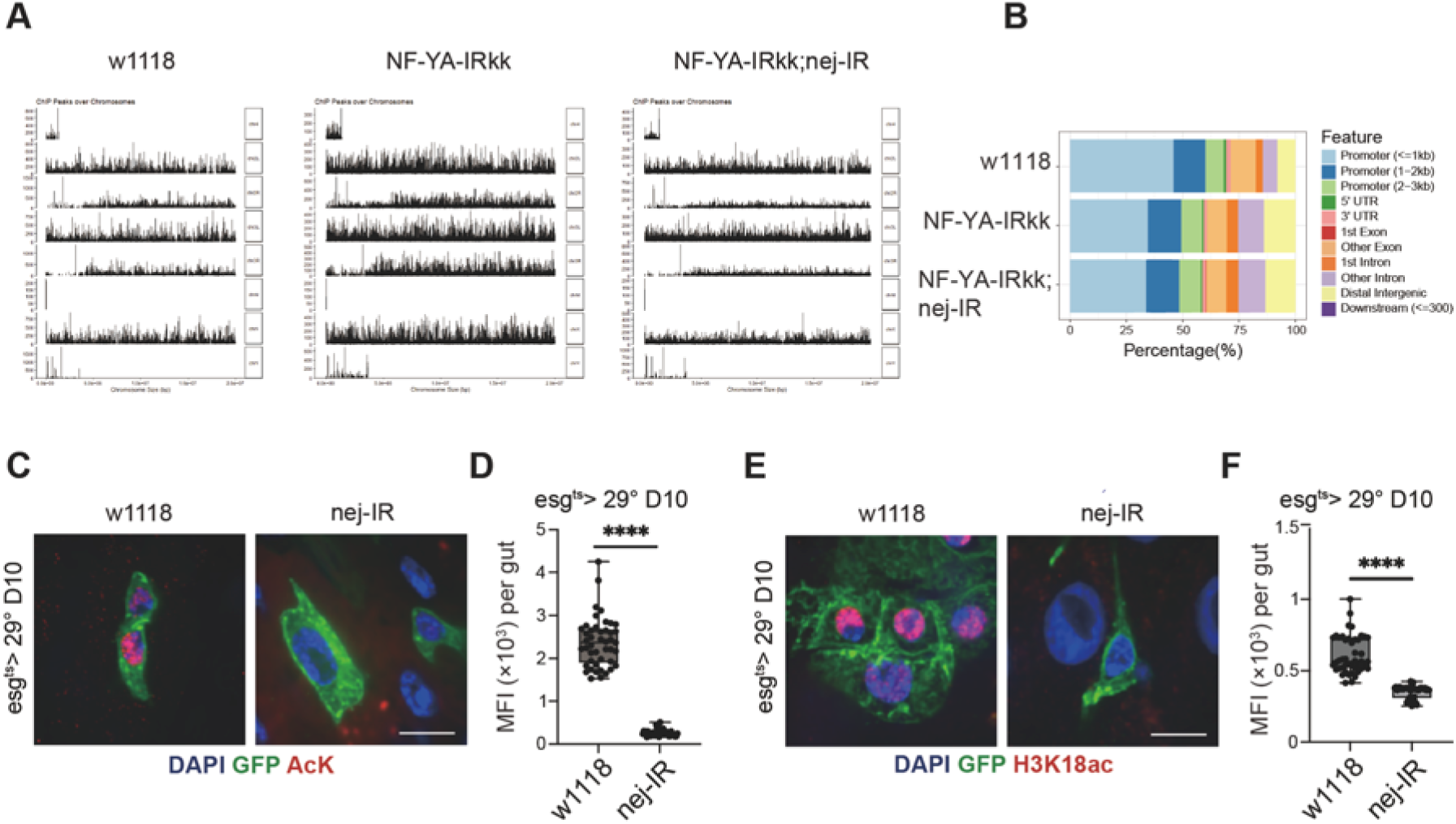
nej is required for basal chromatin acetylation in intestinal progenitor cells. (A) Genome wide distribution of ATAC-seq peaks across chromosomes in the indicated genotypes (w1118, NF-YA-IR, and NF-YA-IR; nej-IR), showing the overall quality and coverage of the ATAC-seq datasets. (B) Genomic feature annotation of ATAC-seq peaks in the indicated genotypes. Peaks are predominantly enriched in promoter-proximal regions. (C and D) Representative images (C) and quantification (D) of pan-acetyl-lysine (AcK) staining in esg+ progenitor cells from control and nej-IR midguts. Blue, DAPI; green, GFP; red, AcK. Scale bar, 10 μm. (E and F) Representative images (E) and quantification (F) of H3K18ac staining in esg+ progenitor cells from control and nej-IR midguts. Blue, DAPI; green, GFP; red, H3K18ac. Scale bar, 10 μm. *(Statistical note for D and F: Data are presented as box-and-whisker plots. Analyzed by two-tailed unpaired Student’s t-test; ***P < 0.0001).

**Figure S6.**
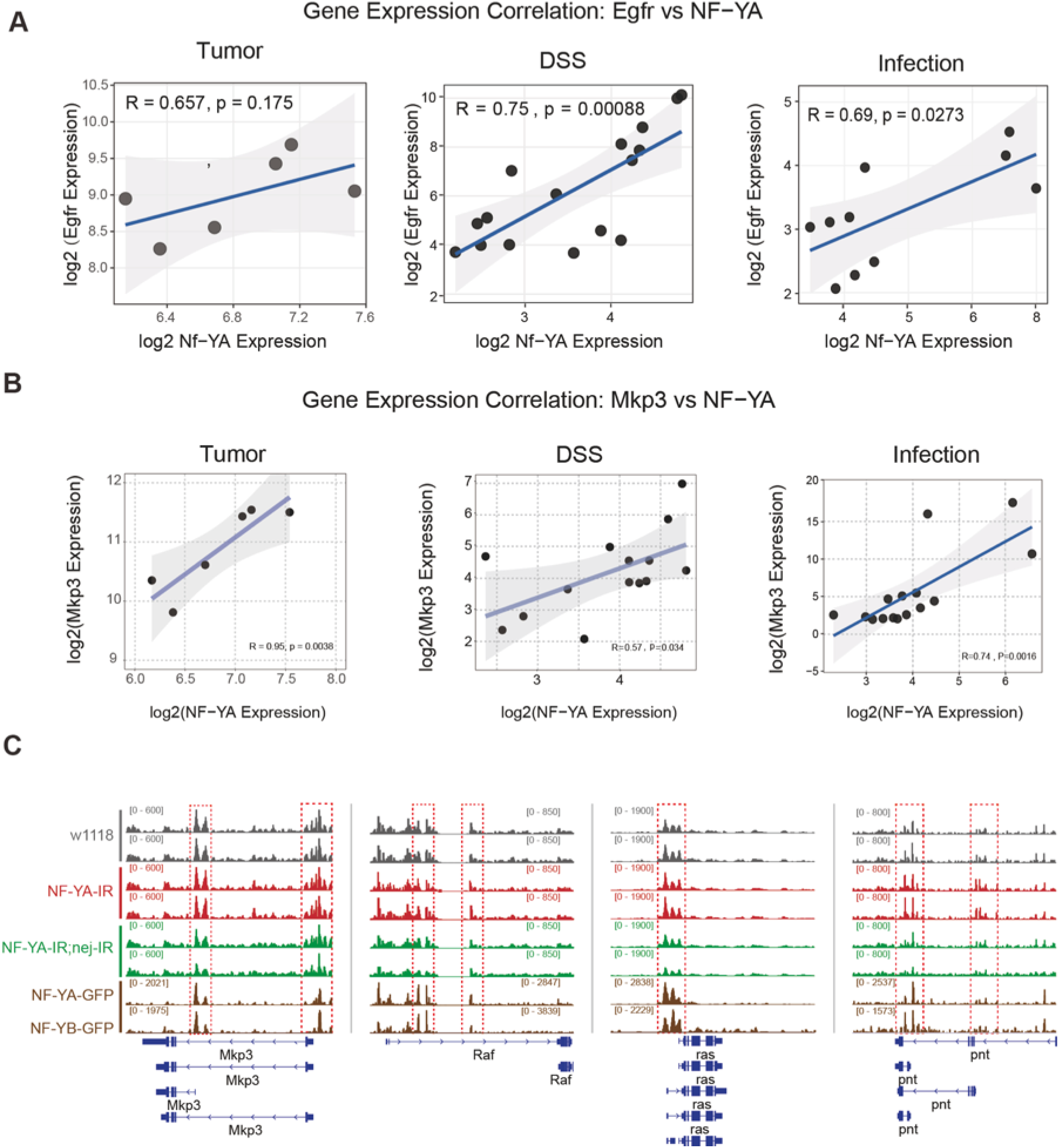
MAPK pathway genes are direct and functionally relevant targets of the NF-Y complex. (A and B) Spearman correlation analyses between the expression levels of NF-YA and EGFR (A), or NF-YA and Mkp3 (B) across published RNA-seq datasets of midgut tumors, DSS injury, and enteric infection (combined *P. entomophila* and *Ecc15* samples). Correlation coefficients (R) and P-values are indicated in the respective panels. (C) Representative genome browser tracks at the Mkp3, Raf, Ras, and pnt loci showing ATAC-seq signals in w1118, NF-YA-IR, and NF-YA-IR; nej-IR midguts, aligned with NF-YA-GFP and NF-YB-GFP CUT&Tag tracks. Red dashed boxes highlight key regulatory regions demonstrating dynamic chromatin accessibility changes linked to NF-Y occupancy.

